# Accurate Predictions of Molecular Properties of Proteins via Graph Neural Networks and Transfer Learning

**DOI:** 10.1101/2024.12.10.627714

**Authors:** Spencer Wozniak, Giacomo Janson, Michael Feig

## Abstract

Machine learning has emerged as a promising approach for predicting molecular properties of proteins, as it addresses limitations of experimental and traditional computational methods. Here, we introduce GSnet, a graph neural network (GNN) trained to predict physicochemical and geometric properties including solvation free energies, diffusion constants, and hydrodynamic radii, based on three-dimensional protein structures. By leveraging transfer learning, pre-trained GSnet embeddings were adapted to predict solvent-accessible surface area (SASA) and residue-specific p*K_a_* values, achieving high accuracy and generalizability. Notably, GSnet outperformed existing protein embeddings for SASA prediction, and a locally charge-aware variant, aLCnet, approached the accuracy of simulation-based and empirical methods for p*K_a_* prediction. Our GNN framework demonstrated robustness across diverse datasets, including intrinsically disordered peptides, and scalability for high-throughput applications. These results highlight the potential of GNN-based embeddings and transfer learning to advance protein structure analysis, providing a foundation for integrating predictive models into proteome-wide studies and structural biology pipelines.

## INTRODUCTION

The three-dimensional (3D) structure of a molecule, typically represented by Cartesian coordinates, contains essential information for deriving its physicochemical and geometric properties^1–3^. Computer programs have long been used to calculate various molecular properties from structures^4–6^, especially properties that are unattainable or impractical to determine experimentally. This is particularly relevant for large biomolecules like proteins, where knowledge of properties like solvation free energy, hydrodynamic radius, solvent-accessible surface area, or p*K_a_* can provide valuable insights into biological function and downstream tasks, like drug design^7, 8^. Traditional methods, such as numerically solving the Poisson-Boltzmann equation to approximate solvation free energy^9^ or running constant pH molecular dynamics (CpHMD) simulations to estimate p*K_a_* values^10^, can prove to be computationally costly. Thus, these traditional methods may be inadequate for high-throughput predictions in the context of modern protein analysis and drug development^8, 11^.

In recent years, the use of data-driven machine learning (ML) methods has emerged as an alternative solution for rapidly and accurately predicting molecular properties from structure. Such methods have been applied to small molecules^12^ and proteins^13^, and they have been trained with experimental and/or simulated reference data^14^. While many of these models have demonstrated significant potential^15–17^, molecular ML methods, especially those designed for structural analysis of proteins, currently face many challenges. For one, developing a predictive model that is both accurate and generalizable requires large and diverse training data sets, but in practice, experimental data is often difficult to obtain, and computing reference data for a large set of molecules is expensive^14^. Moreover, molecular ML models are often designed to predict a narrow set of properties, so they are typically only useful for the specific tasks they were trained on, and predicting new properties generally requires training novel models with new training data sets^18^.

Transfer learning is a potential strategy to address these challenges. In transfer learning, a model first “learns” a latent representation (*i.e.*, an “embedding”) of input features that is optimized to predict target variables for which there is sufficient reference data^19^. Then, the learned embeddings of a “pre-trained” model can be adapted to novel challenges for which training data is sparse^20^, like predicting p*K_a_* values in small molecules^21^. By leveraging insights gained through pre-training, the model may be able to overcome constraints imposed by limited training data in the novel task^22, 23^. “Learning” in this context is similar to latent learning in psychology, where knowledge is acquired without immediate application but becomes apparent through new challenges^24^.

In our work, we construct graph neural network (GNN) models^25^ to produce latent representations of 3D protein structure via pre-training on supervised biomolecular property-prediction tasks. GNNs have previously been shown to be well-suited for molecular data^26–28^, and they have been employed for producing molecular embeddings^13, 29^. However, in previous work, pre-training was either accomplished via self-supervised training to predict relatively simple geometric features, like interatomic distances and bond angles^13^, or via contrastive learning to add higher resolution to coarse-grained protein structures^30^. Different from those approaches, we trained a global structure embedding network (GSnet) and an atomic local charge-aware embedding network (aLCnet) to produce embeddings that capture structural determinants of complex physicochemical features at different levels of resolution.

To obtain large data sets for pre-training, we leveraged ML methods for accurately predicting 3D protein structures from sequence^31, 32^. We primarily used 3D protein models from the AlphaFold Protein Structure Database^33^ and calculated molecular properties using standard physics-based approaches. Specifically, we trained GSnet to predict the radius of gyration (*R*_*g*_), hydrodynamic radius (*R*_*h*_), molecular volume (*V*), translational diffusion constant (*D*_*t*_), rotational diffusion constant (*D*_*r*_), and solvation free energy (Δ*G*_*sol*_) of a protein from its 3D structure. The resultant network was able to predict these target molecular properties with high accuracy, and prediction accuracy remained high when applied to experimental structures from the protein data bank (PDB) and intrinsically disordered peptides (IDPs), even without including such proteins in the training set, demonstrating the broader transferability of our network.

We then applied this pre-trained model to the prediction of molecular solvent-accessible surface area (SASA). Notably, the model was not originally trained to predict SASA, yet it predicted this property with high accuracy by leveraging the learned representations from pre-training. This outcome is notable when compared to other existing protein embedding models, such as GearNet^13^, another GNN that generates structural protein embeddings, and ESM-2^32^, a large language model (LLM) trained on extensive protein sequence databases that has been shown to excel in several structure prediction tasks, including protein structure prediction at atomic resolution. Neither of these alternative embeddings demonstrated similar success in SASA prediction, highlighting a unique advantage of GSnet.

Building on this, we next applied our model to predict p*K_a_* values of amino acid residues within protein structures. Previous studies have utilized representation learning for p*K*_a_ predictions with small molecules^21^ and proteins^34^, but GSnet and aLCnet embeddings are more general as, in principle, their learned representations could be leveraged for predicting other molecular properties. The GSnet and aLCnet models allowed for rapid predictions with similar or better accuracy than previously proposed physics-based and ML-based p*K_a_* predictors^34–36^. Our best predictor approaches an accuracy of 0.9 p*K_a_* units, and since the underlying GNN-based network is relatively lightweight, it is possible to rapidly predict ionization states of amino acids, even in very large complexes, or for large numbers of protein structures appropriate for proteome-scale annotation using experimental or modeled structures as input. We again compared our embeddings with GearNet^13^ and ESM-2^32^, but we were unable to exceed the performance of the null model for p*K_a_* prediction utilizing these other embeddings, demonstrating that GSnet and aLCnet embeddings capture richer information relevant to p*K_a_* prediction than these other methods.

## METHODS

### Datasets

GSnet was pre-trained on the Swiss-Prot subset of the UniProt KnowledgeBase (UniProtKB/Swiss-Prot), utilizing protein structures as predicted by AlphaFold^31,33^. Of the 542,378 proteins in this dataset, we only calculated reference values for a random subset of 153,513 of them due to considerable computational demands. We utilized HYDROPRO^37^ to compute reference values for hydrodynamic radius (*R_H_*), translational diffusion constant (*D_t_*), rotational diffusion constant (*D_r_*), and volume (*V*) using an atomistic representation. Radius of gyration (*R_g_*) was calculated using the MDTraj library in Python^38^ for the same subset of proteins.

We utilized the APBS software suite to compute solvation free energies (Δ*G_sol_*) of proteins from PDB structure according to the linearized Poisson-Boltzmann equation^5^. PQR files, which include charges, were generated via PDB2PQR using the CHARMM c36 forcefield^39^. The grid spacing was set to 0.15 Å, and dimensions were set such that the box was 10% wider than the protein in all three spatial dimensions. Because of the high computational expense of running APBS, Δ*G_sol_* calculations were only obtained for a subset of 30,114 proteins out of the set of 153,513 for which we had values for hydrodynamic properties.

The entire dataset was randomly split into training and validation sets at an approximate ratio of 9:1. The target values in both sets were normalized according to the mean and standard deviation of the data in the training set. Altogether, the training set consisted of 138,290 total structures ranging from 16 to 1,015 residues long, while the validation set consisted of 15,223 total structures ranging from 16 to 966 residues long.

An additional dataset consisting of 123 protein structures was used as a test set. This set was obtained via PISCES^40^ and contained 123 dissimilar (<20% identity), small PDB structures, all of which had a resolution less than 1.5 Å. Another test set consisting of exclusively intrinsically disordered peptides (IDPs) was constructed, which contained ensembles of 100 structures for 45 distinct peptides (4,500 total structures). These ensembles were generated via COCOMO coarse-grained simulations^41^, followed by all-atom reconstruction via cg2all^42^. We also constructed a training set consisting of the ensemble of 100 structures for the shortest IDP, angiotensin, to fine-tune the model for predictions on IDPs.

For training on molecular solvent accessible surface area (SASA), we calculated reference values using the MDTraj library^38^ with atomistic resolution for the same set of 153,513 protein structures described above, with identical splitting into training and validation sets.

For fine-tuning the GSnet to target residue-level SASA (rSASA) values, we calculated reference values with the MDTraj library^38^ for a random subset of 259,049 residues from AlphaFold2 models for sequences in the UniProtKB/Swiss-Prot database^31, 33^, with random splitting into training and validation sets at an approximate ratio of 9:1.

For making p*K_a_* predictions, we utilized the PHMD549 dataset, consisting of p*K_a_* data obtained via constant pH molecular dynamics (CpHMD) simulations, and the EXP67S dataset, consisting of 167 experimental p*K_a_* data points, both of which were proposed by Cai *et al.*^36^. We also constructed our own datasets consisting entirely of experimental p*K_a_* values based on data obtained from PKAD-1^43^ and PKAD-2^44^, as well as from the experimental references listed in Chen *et al*.^35^, Gokcan *et al*.^34^, and Wilson *et al*.^45^. The sources of our data are outlined in **Fig. S1**. All PDB structures were obtained from the Protein Data Bank according to PDB codes provided with the datasets.

Initially, all p*K_a_* data from PKAD-1 were assigned to a set. Non-identical data points outside of PKAD-1 were then compared against all entries in this set. If a data point was structurally similar to (see below) any point in the set, it was added to this set. Otherwise, it was assigned to a separate set. This process produced two non-overlapping datasets: a larger set of 1,932 total points and a smaller set of 237 total points.

The larger set (1,932 points) was split into training and validation sets at an approximate ratio of 9:1, yielding a training set of 1,738 entries and an initial validation set of 194 entries. Within the training set (MSU-pKa-training), structurally similar data points were allowed, as this did not impact model evaluation. However, for the validation set, any similar entries were removed, leaving 52 unique data points in the final validation set (MSU-pKa-validation).

The smaller set (237 points) was used as the basis for an independent test set. To ensure the test set contained only unique data points, we removed any structurally similar entries within this set, resulting in a final test set (MSU-pKa-test) containing 143 unique data points. This process not only ensured uniqueness of each data point in the test set, but it also ensured that the test set remained fully independent of the training and validation sets, preventing data leakage in the construction of the MSU-pKa-test, as seen in other studies and noted by Cai *et al*.^36^. We note that our protocol for preventing data leakage may be more stringent than protocols in similar efforts described in recent papers^46, 47^, where the same residues in proteins with highly similar sequences appear to be considered non-redundant^46^, or where the same residues in the same structures but with different PDB codes are present in published training and test sets^47^.

### Similarity Classification Strategy

To determine structural similarity, 334 total sequences from PKAD-1, PKAD-2, and other sources were clustered using CD-HIT with a sequence identity threshold of 0.5, yielding 71 clusters. We then aligned the sequences within each cluster using ClustalOmega^48^.

A data point (*i.e.*, an ionizable residue) was considered “similar” to another point if all of three conditions were met:

1. The parent sequence was found in the same cluster as the other data point.
2. The residue was in the same alignment column as the residue of the other data point.
3. The residue was of the same amino acid type as the residue of the other data point.

This strategy was used to remove structurally similar entries within the initial validation and test sets. This procedure also ensured that the training set *S*_Train_ and test set *S*_Test_ did not contain any data points corresponding to structurally equivalent residues in proteins with high sequence similarity. Specifically, for each data point *x* ∈ *S*_Train_, we ensure there exists no corresponding data point *y* ∈ *S*_Test_, such that the following conditions hold simultaneously:

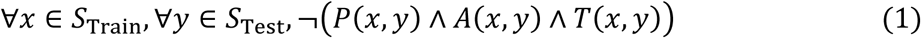

where:

- *P*(*x*, *y*) is true if proteins corresponding to data points *x* and *y* have a sequence identity greater than 0.5,
- *A*(*x*, *y*) is true if the residues corresponding to data points *x* and *y* are in the same alignment column, and
- *T*(*x*, *y*) is true if the residues corresponding to data points *x* and *y* are of the same amino acid type.

We also verified this condition was satisfied between the PHMD549 dataset and the EXP67S dataset, and between the PHMD549 dataset and MSU-pKa-test, to prevent data leakage during transfer learning. We found that there was 1 data point in the PHMD549 dataset, 4O6U_A:134:HIS, that was similar to a data point in MSU-pKa-test, 1B2V_A:133:HIS, so it was not used in pre-training aLCnet.

### Input Features & Graph Construction

An overview of the construction of GSnet is shown in **Fig. 1** and input features are listed in **Table S1**. For each protein in a dataset, an initial graph *G* = (*V*, *E*) was constructed with E(3) invariant features, with nodes *V* representing the residues within the protein and edges *E* between any residues with alpha carbons within 15 Å of each other. The node features *h*_*i*_ ∈ ℝ^*d*^ for each node *i* ∈ *V* incorporated high-dimensional amino acid embeddings, information about dihedral angles, and the distance (in Å) between the constitutive Cα and the protein center of mass. The amino acid embeddings were generated via the torch.nn.Embedding class in PyTorch, and they were learned during model training, with the aim of capturing relevant properties of the amino acids in the context of overall protein structure. The dihedral angle information incorporated the φ and ψ angles, as well as the first three χ angles (where applicable). This dihedral information incorporated both sine and cosine encodings, as well as a mask (0 if the angle does not exist; 1 if the angle exists), totaling 15 values per node (*i.e.*, residue). We included dihedral angles as node features to provide the model with information about the spatial configuration of a protein’s backbone and side chains, which should effectively provide the model with higher resolution. The distance (in Å) between the constitutive Cα and center of mass of the protein was used as a node feature with the aim of providing the model information about each residue’s location within the overall protein structure. Multilayer perceptrons (MLPs) were trained to learn high-dimensional tensor representations of non-discrete input data, including the dihedral information and distances, and they were also employed for dimensionality reduction of input node features after concatenation (see **Fig. S2**). Edges *E* of the graph were generated between any residues that were within 15 Å of each other, with distances *d*_*i*_ (in Å) calculated between the Cα atoms of residues *i* and *j* that represented an edge’s corresponding nodes:

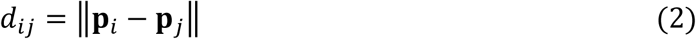

where *p*_*i*_ and *p*_*i*_ are the Cartesian coordinates of the Cα atoms of residues *i* and *j*, respectively. We then applied Gaussian smearing to transform these scalar distances into higher-dimensional edge features *e*_*ij*_ according to:

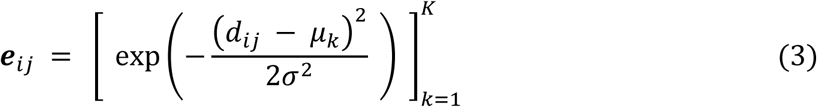

where μ_k_ are the centers of linearly spaced Gaussian basis functions, σ is the spacing between consecutive μ_k_ values, and *K* is the total number of Gaussians used (*i.e.*, the dimensionality of the edge features). We set *K* = 300, with μ_k_ linearly spaced from 0.0 to 15.0 Å in increments of approximately 0.05017 Å. Thus, the resulting edge features *e*_*i*_ are 300-dimensional vectors. An overview of the extraction of GSnet node and edge embeddings from input features is shown in **Fig. S2**.

**Figure 1.**
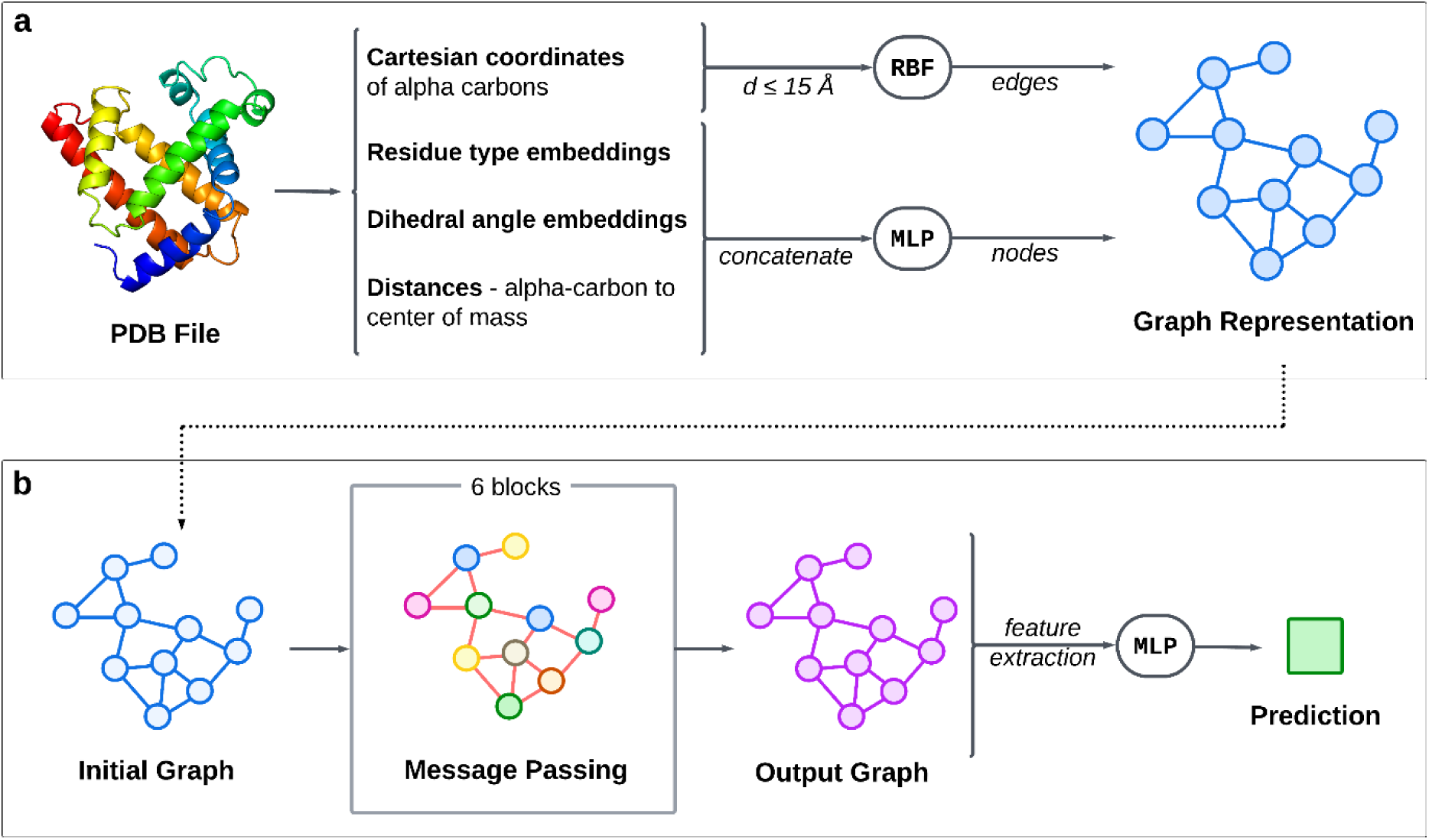
Overview of GSnet architecture. **(a)** Graph construction. Input node and edge features are extracted from the PDB structure to create a graph representation of the protein via multilayer perceptrons (MLPs) and a radial basis function (RBF) (see **Fig. S2** for more detail). **(b) Neural network architecture.** This initial graph is passed to a GNN consisting of six message-passing blocks (see **Fig. S4** for more detail), which ultimately generates an output graph with a high-dimensional embedding for each node. Features are extracted from the output graph and passed to a multilayer perceptron (MLP) to make final predictions.

An analogous graph construction was performed for aLCnet, shown in **Fig. 2** with input features listed in **Table S2**. When using aLCnet, an initial graph *G* = (*V*, *E*) was also constructed with E(3) invariant features, but with nodes *V* for each carbon, hydrogen, nitrogen, oxygen, and sulfur atom within a 10 Å radius surrounding and including the alpha-carbon of the residue of interest, rather than for every residue in an entire protein. Here, node features *h*_*i*_ ∈ ℝ^*d*^ for each node *i* ∈ *V* incorporated the same amino acid embeddings as GSnet, along with similar embeddings for atom type, as well as atomic partial charge as determined via PDB2PQR^5^. Residue and atom embeddings were included with similar reasoning to GSnet residue embeddings. Atomic charge was incorporated to provide the model critical information about electrostatic interactions, which directly influence p*K_a_*. Like with GSnet, MLPs were trained to learn high-dimensional tensor representations of non-discrete input data (partial charges in this case), and for dimensionality reduction of input node features after concatenation (see **Fig. S3**). Edges *E* were generated between any atoms that were within 5 Å of each other, with the only edge feature *e*_*i*_ again derived according to **Eqs. 2** and **3**, but where *p*_*i*_ and *p*_*i*_ are the Cartesian coordinates of atom *i* and *j*, respectively, and with μ_k_ linearly spaced from 0.0 to 5.0 Å in increments of approximately 0.0167 Å.

**Figure 2.**
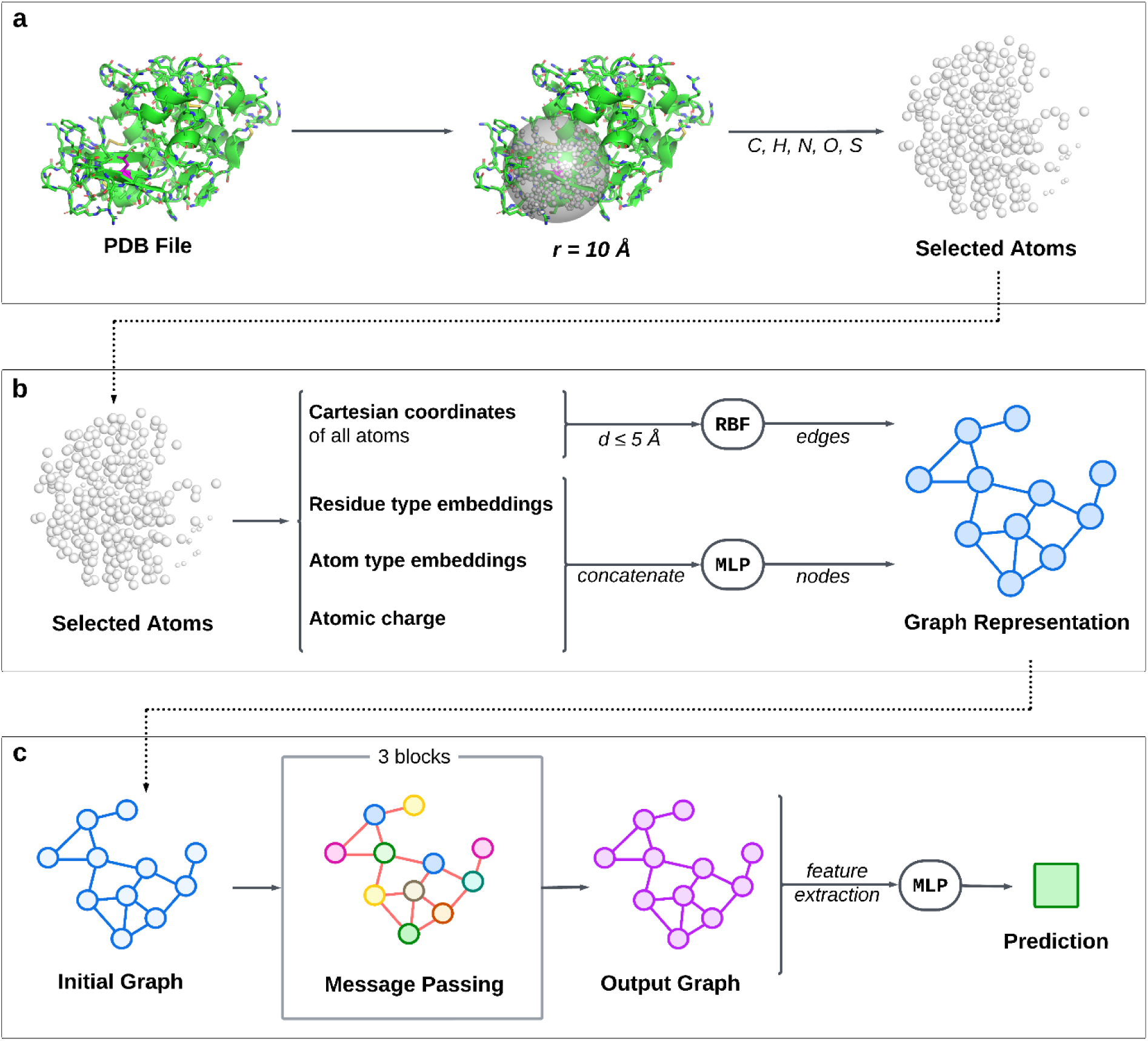
Overview of aLCnet architecture. **(a)** Atom selection. Carbon, hydrogen, nitrogen, oxygen, and sulfur atoms within ten angstroms of the alpha carbon of the residue of interest are selected from the original PDB structure. **(b) Graph construction.** Input node and edge features are extracted from the selected atoms to create a graph representation of the protein environment via a multilayer perceptron (MLP) and radial basis function (RBF), respectively (see **Fig. S3** for more detail). **(c) Neural network architecture.** Like GSnet, the initial graph is passed to a GNN, this one consisting of three message-passing layers (see **Fig. S4** for more detail), which ultimately generates an output graph with a high-dimensional embedding for each node. Features are extracted from the output graph and passed to a multilayer perceptron (MLP) to make a final prediction.

### Neural Network Architecture

With GSnet, the initial protein graph *G* = (*V*, *E*) with 150-dimensional node features *h*_*i*_ ∈ ℝ^1^^50^ for each residue *i* ∈ *V* is passed through six custom transformer message passing layers based on those described by Shi *et al.*^49^. Each message-passing layer applies attention to update the node features by attending over the neighboring nodes in *G*.

Before message passing, edge features *e*_*i*_ are transformed via:

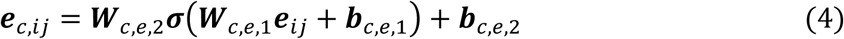

where σ is the shifted softplus activation function defined as σ(*x*) = ln(1 + e^*x*^) − ln(2)1, and 300-dimensional edge features *e*_*i*_ to 150-dimensional edge features *e*_c,*ij*_, in congruence with the dimensionality of the node features. Note that the subscript *c* denotes the attention head index, but GSnet only employs one attention head.

Next, the node features 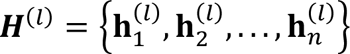 in the graph at layer *l* are updated through the attention-based message-passing mechanism described by Shi *et al.*^49^. These updated node features 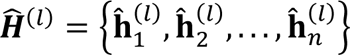 were then passed through an additional residual connection to arrive at the updated node features 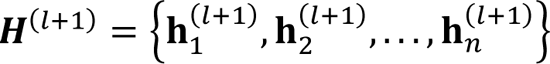 at the next layer *l* + 1 according to:

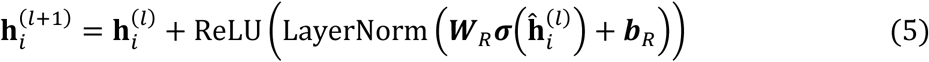

where LayerNorm is the layer normalization operation described by Ba *et al.*^50^. **Fig. S4** shows a diagram outlining the entire message passing process.

We evaluated different message passing architectures, namely Schnet^51^, EGNN^52^, and TransformerConv^49, 53^, as well as various numbers of attention heads, layers, and hidden channels, and the best performance we could achieve was done with these layers and parameters.

For global property predictions, the element-wise mean μ across embeddings *h*_*i*_ for all nodes *i* ∈ *V*, following message-passing, was computed according to:

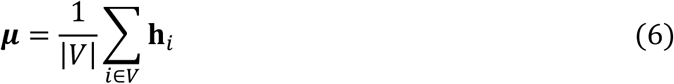

where |*V*| is the total number of nodes in the graph. This global mean was then passed to an output MLP consisting of four linear layers with 1024 hidden channels each, and three shifted softplus (SSP) activation layers. This output MLP was trained to output the six target values, or for molecular SASA, the one target value.

To make residue-level SASA predictions, an architecturally identical output MLP was used as with molecular SASA; however, the 150-dimensional node embedding specific to the residue for which a prediction was being made *h*_*i*_ was passed to the MLP, rather than the component-wise mean μ (see **Table S3**). To make per-residue p*K_a_* predictions, a total of seven 150-dimensional features were extracted and concatenated, resulting in a 1,050-dimesnsional feature (see **Eqs. 6-8, Table S3**). These features included μ and *h*_*i*_, as well as five element-wise means across nodes representing residues within a spatial radius *r* (in Å) from the residue of interest μ_*r*_:

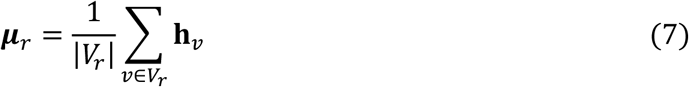

where *V*_*r*_ ⊆ *V* denotes the subset of nodes corresponding to residues within *r* of residue of interest and |*V*_*r*_| is the number of nodes in this subset. Specifically, we used radii 6, 8, 10, 12, and 15 Å.

The resulting tensor was then passed to an output MLP which consisted of six linear layers with 1024 hidden channels, six dropout layers with 20% dropout, and five SSP activation layers.

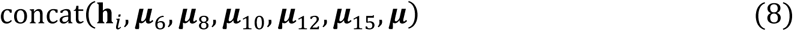

The architecture for aLCnet followed a similar pattern as GSnet, but with 75-dimensional node features *h*_*i*_∈ ℝ^75^ for each atom in the selection and only three of the previously described transformer-based message-passing layers (see **Fig. S4**), each with three attention heads.

For p*K_a_* predictions, three 75-dimensional features were extracted and concatenated, namely the element-wise mean across all atoms in the selection μ (calculated via **Eq. 6**), the embedding representing the Cα atom in the residue of interest *h*_cα_, and the element-wise mean across nodes representing atoms in the residue of interest μ_*aa*_ (calculated via **Eq. 9**):

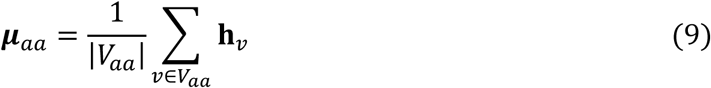

where *V*_*aa*_ ⊆ *V* denotes the subset of nodes corresponding to atoms that compose the residue of interest and |*V*_*aa*_| is the number of nodes in this subset. These features were concatenated according to **Eq. 10:**

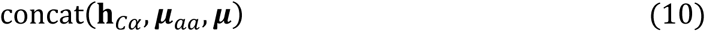

The output MLP here was identical to the one used for p*K_a_* predictions with GSnet, with the only difference being a 225-dimensional, rather than 1,050-dimensional, input.

### Training

To train GSnet on the geometric properties, as well as Δ*G_sol_* (for which reference values were only available for 30,114 out of 153,513 proteins in the training set), a mask was applied to set the gradients to zero for instances where Δ*G_sol_* reference values were not available, ensuring that model weights remained unchanged by missing values while still being updated based on available target values^54^.

We trained 20 models for each type of GNN layer, totaling 60 models, using the PyTorch^55^ and PyTorch Geometric libraries^56^. We utilized the Adam optimizer^57^ with an initial learning rate of 1E-4, adjusted to 1E-5 after 50 epochs by a scheduler. Models were trained to minimize the mean squared error (MSE) loss across the six target values with a batch size of 64. Training was terminated when there was no improvement in validation performance for ten consecutive epochs (**Fig. S5**). The model with the lowest validation MSE loss (of 20) across the six targets was selected for further evaluation on the test sets and for transfer learning applications.

To fine-tune GSnet for intrinsically disordered peptide (IDP) predictions, we first loaded the weights and biases of the GNN as obtained via pre-training on the original six target values. We then trained from here in the same fashion as the original training runs, using the ensemble of 100 angiotensin structures as the training set.

GSnet was fine-tuned similarly for molecular SASA predictions, but the parameters of the pre-trained GNN were kept fixed such that only the output MLP was trained. This was done to test the broader transferability of the original weights and biases.

For residue-level SASA predictions, we trained models using two approaches. In the first, we applied the original GSnet (as trained on the six original properties) with fixed weights and only trained an output MLP. In the second, we trained an output MLP, as well as the fine-tuned GSnet itself by allowing optimization of the GNN weights based on the residue-level SASA dataset. Training on this dataset was performed similarly to before, but training was terminated after ten epochs. In each approach, five models were trained from the same original parameters, and the “best” model was selected based on lowest root mean squared error (RMSE) on the validation set.

For p*K_a_* predictions based on GSnet, we used four training approaches: the initial GSnet with fixed GNN weights, the initial GSnet with GNN weights allowed to be optimized, the fine-tuned GSnet for predicting residue-level SASA values with fixed GNN weights, and the fine-tuned GSnet for predicting residue-level SASA values with GNN weights allowed to be optimized. In all cases, an output MLP was trained to map GNN embeddings to predictions. For each approach, we trained 20 total models to minimize MSE loss across the training p*K_a_* dataset, utilizing the Adam optimizer with a learning rate of 1E-4 and a batch size of 64. A 20% dropout was necessary to prevent overfitting and to optimize validation performance. These models were trained for 100 epochs, and their performance was evaluated based on RMSE on the validation set. The ten best performing models based on validation RMSE were further evaluated on the test set to generate statistics. This procedure was conducted for both the datasets proposed by Cai *et al*.^36^ and for our datasets. An overview of the entire training process for GSnet is shown in **Fig. S6**.

With aLCnet, we also utilized transfer learning for p*K_a_* prediction, but we did not pre-train with the same data as with GSnet. Instead, we first trained 20 randomly initialized aLCnet models on the simulated data contained in the PHMD549 dataset. The top 10 models based on validation RMSE were selected and evaluated on the EXP67S test set. We then selected the pre-trained model with the lowest validation RMSE for fine-tuning using MSU-pKa-training and MSU-pKa-validation. Again, 20 training runs were performed and the top ten models were evaluated on MSU-pKa-test. An overview of the entire training process for aLCnet is shown in **Fig. S7**.

### Alternative Embeddings

When training our GNN models on the original six values, molecular SASA, and p*K_a_*, we compared their performance to the performance of ESM-2^32^, a large language model, and GearNet^13^, another structure-based GNN. The 650 million parameter ESM model that we used generated 1,280-dimensional embeddings for each residue, while the GearNet model generated 3,072-dimensional embeddings for each residue. For global property predictions with these models, the mean of embeddings was calculated as in **Eq. 6** and used as an input feature. For p*K_a_* predictions, model embeddings were concatenated in the same fashion as GSnet embeddings (see **Eqs. 6-8, Table S3**) to obtain input features. These features were passed to identical output MLPs as were used in the different applications of our model, with the only difference being the number of input channels, based on the embedding dimension of the given model.

Code is available on GitHub at https://github.com/feiglab/ProteinStructureEmbedding

## RESULTS AND DISCUSSION

### Global structural embeddings

To create global structure embeddings, a GNN and output MLP (Fig. 1) were trained with the goal of simultaneously predicting six molecular properties of proteins, namely free energy of solvation (Δ*G_sol_*), radius of gyration (*R_g_*), hydrodynamic radius (*R_h_*), translational diffusion coefficient (*D_t_*), rotational diffusion coefficient (*D_r_*), and molecular volume (*V*). These properties were selected based on their significance in characterizing protein structure and function: *R_g_* is a geometric property that captures the distribution of atoms within the molecular cavity; hydrodynamic properties like *R_h_*, *D_t_*, and *D_r_* capture the external shape of the protein and its interactions with the surrounding solvent; and Δ*G_sol_* captures information about the charge distribution, allowing the inference of details about electrostatic interactions. The computational calculation of hydrodynamic properties and Δ*G_sol_* via traditional software is very costly, taking up to days for a single protein, highlighting the need for an efficient and accurate ML model.

Various GNN implementations were tested, namely SchNet^51^, an EGNN^52^, and a transformer^49, 53^, to determine which architecture provides the highest predictive accuracy for these properties. We found similarly high prediction accuracy across all three of these GNN architectures when applied to *R_g_*, *R_h_*, *D_t_*, *D_r_*, and *V* (**Table 1**), and validation set performance was only moderately worse than training set performance (**Table S4**). Among our GNN models, the transformer architecture performed slightly better, achieving the lowest mean absolute percent errors (MAPEs) and root mean squared errors (RMSEs) across the target values (**Table 1**). Only for Δ*G_sol_*, the EGNN model was slightly better (**Table 1**). Therefore, we adopted the transformer model (referred to as GSnet) for transfer learning applications.

**Table 1.** Validation set performance for prediction of global molecular properties. Mean absolute percent errors (MAPE) and root mean square errors (RMSE) are reported as the mean and standard errors from 20 independent training runs (except for ESM-2), with the value for the best overall model (chosen by lowest MSE across all target values) given in parentheses. Bold values indicate the best performance across all models for a given target value.

| | Model | $\Delta G_{\text{sol}}$<br>[kJ/mol] | $R_g$ [Å] | $R_h$ [Å] | $D_t$<br>[nm<sup>2</sup>/μs] | $D_r$ [μs<sup>-1</sup>] | $V$ [nm<sup>3</sup>] |
| --- | --- | --- | --- | --- | --- | --- | --- |
| **MAPE (%)** | *Transformer (GSnet)* | **4.21 ± 0.04**<br>**(3.89)** | **0.97 ± 0.01**<br>**(0.82)** | **0.67 ± 0.00**<br>**(0.65)** | **0.65 ± 0.00**<br>**(0.65)** | **2.79 ± 0.03**<br>**(2.47)** | **1.10 ± 0.01**<br>**(1.12)** |
|  | *EGNN* | **4.15 ± 0.03**<br>(3.95) | 1.09 ± 0.01<br>(1.22) | 0.69 ± 0.01<br>(0.73) | 0.68 ± 0.00<br>(0.71) | 2.93 ± 0.04<br>(2.71) | 1.13 ± 0.01<br>(1.20) |
|  | *SchNet* | 5.71 ± 0.03<br>(5.84) | 1.25 ± 0.02<br>(1.15) | 0.81 ± 0.00<br>(0.80) | 0.78 ± 0.00<br>(0.80) | 3.41 ± 0.03<br>(3.60) | **1.11 ± 0.01**<br>**(1.12)** |
|  | *GearNet* | 6.75 ± 0.03<br>(6.67) | 3.05 ± 0.01<br>(3.00) | 1.97 ± 0.00<br>(1.96) | 1.97 ± 0.00<br>(1.95) | 6.64 ± 0.02<br>(6.66) | 4.07 ± 0.01<br>(4.07) |
|  | *ESM-2<sup>a</sup>* | 9.04<br>(9.04) | 5.02<br>(5.02) | 2.91<br>(2.91) | 2.91<br>(2.91) | 10.2<br>(10.2) | 5.43<br>(5.43) |
| **RMSE** | *Transformer (GSnet)* | 565.5 ± 2.4<br>**(532.0)** | **0.38 ± 0.00**<br>**(0.35)** | **0.31 ± 0.00**<br>**(0.30)** | **0.71 ± 0.00**<br>**(0.71)** | **0.38 ± 0.00**<br>**(0.37)** | **0.66 ± 0.01**<br>(0.65) |
|  | *EGNN* | **555.6 ± 2.8**<br>(538.0) | 0.42 ± 0.00<br>(0.42) | 0.32 ± 0.00<br>(0.31) | 0.73 ± 0.00<br>(0.77) | 0.40 ± 0.00<br>(0.39) | **0.66 ± 0.01**<br>**(0.64)** |
|  | *SchNet* | 844.7 ± 2.1<br>(837.0) | 0.47 ± 0.00<br>(0.46) | 0.37 ± 0.00<br>(0.36) | 0.84 ± 0.00<br>(0.86) | 0.48 ± 0.00<br>(0.49) | **0.67 ± 0.01**<br>(0.71) |
|  | *GearNet* | 1031.1 ± 3.0<br>(1032.4) | 1.65 ± 0.00<br>(1.62) | 1.04 ± 0.00<br>(1.03) | 2.20 ± 0.00<br>(2.18) | 0.89 ± 0.00<br>(0.88) | 3.51 ± 0.01<br>(3.58) |
|  | *ESM-2<sup>a</sup>* | 1298.0<br>(1298.0) | 2.24<br>(2.24) | 1.38<br>(1.38) | 3.29<br>(3.29) | 1.56<br>(1.56) | 3.77<br>(3.77) |

<sup>a</sup>Only one model was trained with ESM-2 due to computational cost.

We also tested alternative pre-trained ML models that generate protein representations, namely ESM-2^32^ and GearNet^13^. An architecturally identical output MLP (except for number of input dimensions, as dictated by the embedding size of the tested model) as used with our GNN models was trained to predict the six properties based on these alternative input embeddings. All our GNN models significantly outperformed networks based on GearNet and ESM-2 embeddings (**Table 1**).

The performance of GSNet for predicting global molecular properties is illustrated further in Fig. 3. Strong correlation is found for the systems in the validation set across the entire range of values, only slightly worse than the correlation obtained for the training set (**Fig. S8**). To confirm that the validation accuracy of the model did not result from data leakage due to homologous structures in the training set, we performed BLAST alignments^58^ for each validation set protein against all proteins in the training set and established a threshold for homology at an E-value^59^ of 10^-5^. We found that the performance of the models on the validation set proteins that did not exhibit homology to any proteins in the training set was roughly the same as for the proteins that did, indicating that the accuracy was independent of data leakage (Fig. 3).

**Figure 3.**
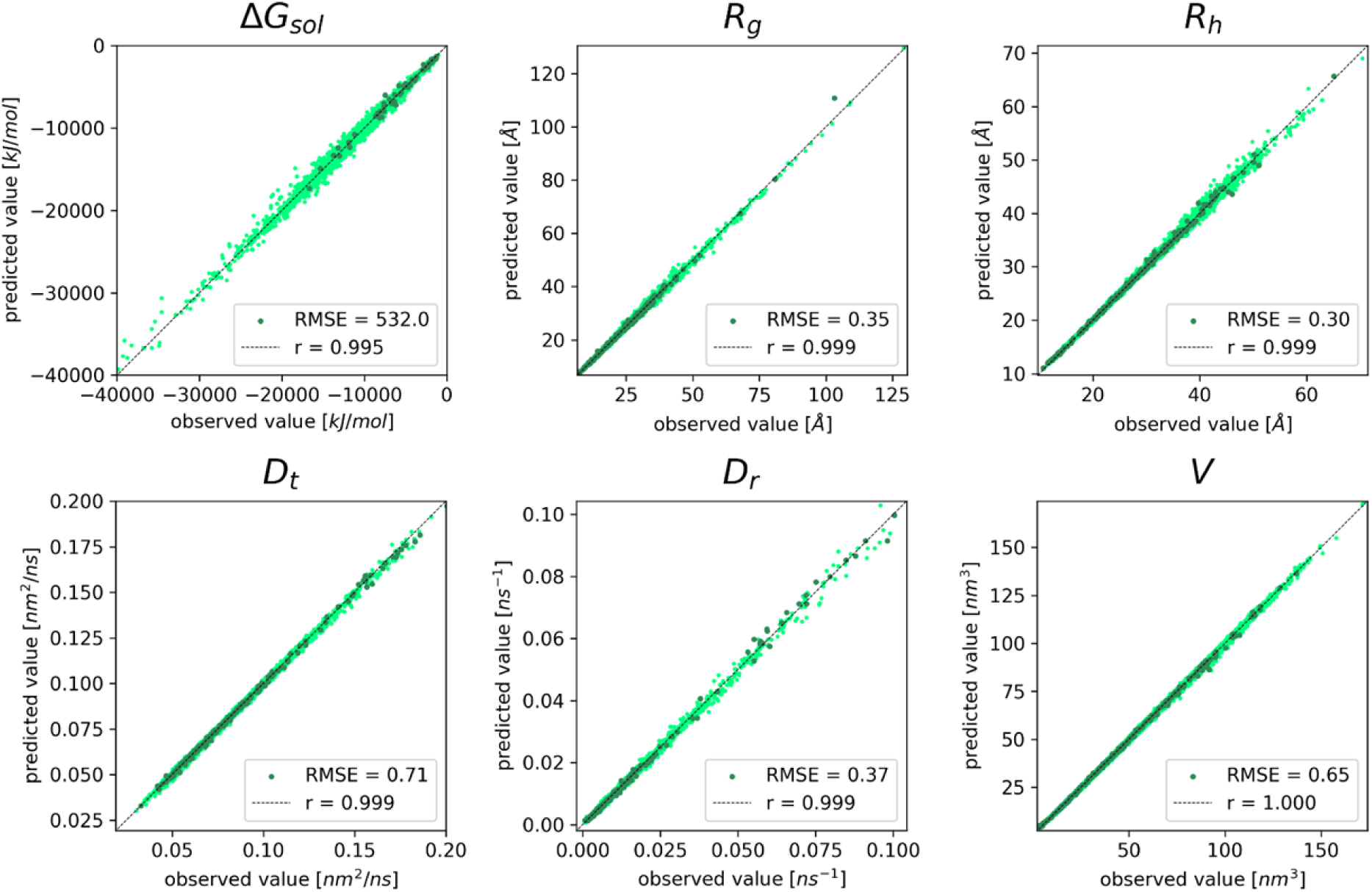
GSnet predictions of global molecular properties vs. reference values. Results are shown for Δ*G_sol_*, *R_g_*, *R_h_*, *D_t_*, *D_r_*, and *V*. Darker points represent structures that are not homologous to any structures in the training set. Pearson correlation coefficients and RMSEs encompass all data shown in the plots. RMSE values are given in units as displayed on the axes of each subplot.

Across all geometric properties, GSnet exhibits an error ranging from 0.65-2.47%, which is lower than HYDROPRO’s error of approximately 4% relative to experimental values^6^. This suggests that, given enough accurate experimental training data, GSnet could surpass HYDROPRO’s accuracy in predicting hydrodynamic observables. However, predictions of Δ*G_sol_*, with an error of 3.89%, are significantly less accurate than continuum electrostatics calculations, where the typical error may be 0.25% or lower^60, 61^. The relatively poor performance of GSnet for predicting Δ*G_sol_* may result from limited training data relative to the other features, or because it is inherently more difficult to predict solvation free energies from structural embeddings alone.

Since GSnet was trained using AlphaFold (AF) models, we further evaluated GSnet on datasets consisting of 123 structures of smaller proteins from the Protein Data Bank. Performance on this dataset was similar to the performance of the model on the larger dataset for *R_g_*, *R_h_*, *D_t_*, *D_r_*, and *V* (**Fig. S9**). However, the model seemed to overestimate on Δ*G_sol_* predictions relative to APBS. Further evaluation of GSnet on a larger set of 610 PDB structures^60^ showed the same tendency (**Fig. S10**) across all PDB structures. To test whether this shift was due to issues with PDB structures, a brief energy minimization was conducted via CHARMM^62^ and reference values were calculated again. On the minimized structures, GSnet underestimated solvation free energies relative to APBS (**Fig. S10**). Subsequently, AF structure predictions for the same subset of proteins were obtained and reference values were calculated for these structures. Interestingly, the model predictions on these structures restored the initial accuracy reported for the validation set without systematic deviations in either direction. This suggests that GSnet learned specific features of AF models when making Δ*G_sol_* predictions. While this may not be a significant issue for the purpose of generating a reusable structural embedding in this work, future efforts may consider how to make predictions of solvation free energies with model-independent accuracies.

Finally, another dataset was considered to explore the transferability of GSnet towards intrinsically disordered peptides (IDPs). IDPs were generally not present in the training set, although some of the training set proteins certainly contained short, disordered elements such as loops as part of folded structures. The IDP dataset consisted of conformational ensembles generated via coarse-grained simulations using COCOMO^41^, followed by all-atom reconstruction^63^. All reference values were obtained in the same manner as for the other datasets. GSnet predicted the target properties again with high correlation for this dataset; however, performance was generally worse compared to folded proteins (**Fig. S11**). The model exhibited a propensity to underestimate *R_g_*, *R_h_*, and *V*, while overestimating *D_t_* and *D_r_* (**Fig. S11**). These shifts are likely due to the decreased compactness of IDPs as compared to folded proteins. Because radius of gyration, hydrodynamic radius, and volume inherently measure a protein’s size and spatial occupancy, GSnet trained on folded proteins may be biased towards predicting lower values, similar to those observed in more compact, folded proteins. Accordingly, considering that diffusion coefficients are inversely related to protein size^64^, the model may anticipate greater values. Additionally, the model did not perform well on very small IDPs, notably angiotensin (8 residues), YESG2 (8 residues), and YESG6 (12 residues), all of which were shorter than any of the proteins found in the original training set. The performance of GSnet on the IDP set shows limitations of GSnet when transferred to structures unlike those in the training set.

To test whether additional training could improve the model, GSnet was fine-tuned with an additional ensemble of 100 angiotensin structures. In the fine-tuned model, most of the systematic shifts were removed and the large errors for very short peptides were eliminated (**Fig. S12**). This indicates that modest data augmentation may be enough to adapt GSnet to structures outside the domain of the original training set.

### Predicting of Global Properties via Transfer Learning from GSnet

To explore the transferability of GSnet embeddings to other global properties, we focused on the prediction of solvent-accessible surface areas (SASA). The parameters of the pre-trained GNN were loaded and kept fixed during this process, while a new, trainable output MLP was introduced to map the element-wise mean of the GNN node embeddings for a given protein structure to its SASA value. After training the new MLP to predict SASA based on the pre-trained GNN embeddings, on the same structures as in the original training set, the model showed excellent performance on the validation data (Fig. 4). For comparison, we also considered the use of embeddings from ESM-2^32^ and GearNet^13^. The training and validation performance were significantly worse when using either of those embeddings (Fig. 4, **Fig. S13**), indicating that GSnet may be a better platform for transfer leaning applications for the prediction of protein structure properties.

**Figure 4.**
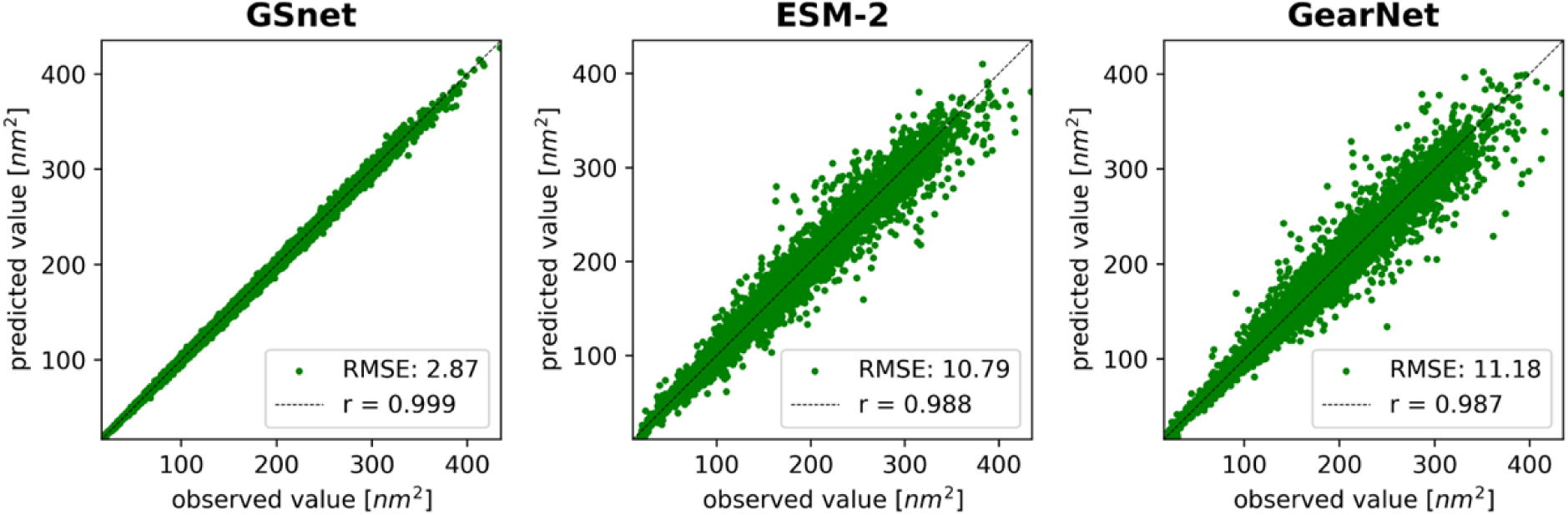
Validation set performance of embedding-based SASA predictors. Correlation coefficients from linear fits are shown in the upper left corner of each plot. RMSE values are given in nm^2^.

Interestingly, the validation set performance of GSnet-based SASA predictions is very similar to the training set performance (**Fig. S13**). This is not the case for GearNet-based SASA predictions where validation set performance is significantly decreased over the training set performance, indicating better transferability with GSnet embeddings. To further explore the transferability of GSnet embeddings, we evaluated the model on three subsets of the original training set with proteins shorter than 50, 100, or 200 residues, respectively. The motivation for considering these subsets was to test whether transferability to larger proteins could be achieved without including such proteins during training, since the calculation of reference values needed for training becomes increasingly costly as the protein size increases. In addition to strictly limiting protein size in the training set, we also considered a data augmentation strategy where a few larger proteins were added to a training set that was otherwise limited in size. Training on size-limited training sets, we found that the accuracy of predictions for proteins larger than the training set limit quickly deteriorated (**Fig. S14**), but the performance was better with augmented training data (**Fig. S15**). Specifically, training on proteins with up to 200 residues augmented by a few larger proteins resulted in fairly accurate validation set predictions (r = 0.997, RMSE = 6 nm^2^) across the entire range of proteins up to 1000 residues (**Fig. S15**). This suggests that transfer leaning based on GSnet does require that the training set domain covers the target application domain, but training of accurate models may be possible with sparser data than what was used in the original training of GSnet.

### Predicting of Local Properties via Transfer Learning from GSnet

We also tested whether GSnet embeddings were useful in making residue-level predictions. We first applied the embeddings of GSnet to predict residue-level SASA values by training an output MLP with fixed, pre-trained GNN parameters. We found relatively poor correlation (r = 0.751) and RMSE (0.36 nm^2^), and the relative errors were significantly larger than for the predictions of the global structures (see **Fig. S16A**). We then tested whether fine-tuning the GNN itself (*i.e.*, allowing the parameters of the pre-trained GNN to be optimized along with the output MLP) can improve accuracy. We found significant improvements in correlation (r = 0.956) and RMSE (0.16 nm^2^) relative to fixed GNN parameters, but the relative errors remained larger than for the prediction of global properties (**Fig. S16B**), indicating that the GSnet embedding may not have enough capacity to predict residue-level properties with the same accuracy as global structural properties.

We then considered the prediction of residue-specific p*K_a_* values. We first focused on using simulated p*K_a_* data from the PHMD549 dataset for training^36^, and then we evaluated the model on the EXP67S test set consisting of experimental p*K_a_* values as in previous work^36^. We used simulated p*K_a_* for training here to have enough training data to evaluate different models and training protocols, whereas testing the resulting models against experimental p*K_a_* values allowed us to compare with other approaches from other groups. Here, we applied four training approaches: the initial GSnet with either fixed or optimized GNN weights, and the fine-tuned GSnet for residue-level SASA predictions, again with either fixed or optimized GNN weights.

As shown in **Table 2**, fine-tuning the model for residue-level SASA predictions before p*K_a_* training resulted in more accurate p*K_a_* predictions, with average RMSE decreasing from 1.39 to 1.31 when not optimizing GNN weights, and decreasing from 1.33 to 1.29 when GNN weights were allowed to optimize. This improvement suggests that fine-tuning on residue-level SASA predictions helped the model better capture local structural properties relevant to residue-specific properties, such as p*K_a_*. Moreover, allowing the GNN to optimize during p*K_a_* training led to marginal, but significant, improvements in accuracy, with average RMSE dropping from 1.39 to 1.33 with the original GSnet weights, and dropping from 1.31 to 1.29 with the fine-tuned GSnet weights on residue-level SASA. This improvement indicates that allowing GNN weights to optimize enables the model to refine its representations further to account for structural features uniquely relevant to p*K_a_* prediction.

**Table 2.** Prediction of experimental p*K_a_* values from the EXP67S test set. Performance with GSnet-based models is given as mean ± SEM with the best test RMSE in parentheses. Performance data for the null model, PROPKA, CpHMD, and PKAI+ was taken from from Cai *et al*.^36^. Performance data for DeepKa was obtained via predictions made with the DeepKa web server^65^.

| Method | RMSE | R | m |
| --- | --- | --- | --- |
| Null Model<sup>a</sup> | 1.44 | - | 0 |
| PROPKA<sup>a</sup> | 1.12 | 0.63 | 0.45 |
| CpHMD<sup>a</sup> | 1.01 | 0.73 | 0.65 |
| PKAI+<sup>a</sup> | 1.30 | 0.45 | 0.15 |
| DeepKa<sup>a</sup> | 0.97 | 0.74 | 0.50 |
| GSnet<sup>b</sup> | 1.39 ± 0.01 (1.37) | 0.40 ± 0.01 (0.42) | 0.14 ± 0.00 (0.15) |
| GSnet (rSASA)<sup>c</sup> | 1.31 ± 0.01 (1.28) | 0.52 ± 0.00 (0.53) | 0.19 ± 0.00 (0.21) |
| GSnet (opt)<sup>d</sup> | 1.33 ± 0.00 (1.31) | 0.49 ± 0.01 (0.52) | 0.19 ± 0.01 (0.23) |
| GSnet (rSASA-opt)<sup>e</sup> | 1.29 ± 0.00 (1.26) | 0.52 ± 0.00 (0.54) | 0.24 ± 0.01 (0.28) |
| aLCnet<sup>f</sup> | 1.10 ± 0.01 (1.08) | 0.63 ± 0.01 (0.67) | 0.38 ± 0.01 (0.48) |

<sup>a</sup>values taken from Cai *et al.*<sup>36</sup>.
<sup>b</sup>using pre-trained GSnet model with fixed GNN weights
<sup>c</sup>using pre-trained GSnet model fine-tuned for residue-level SASA with fixed GNN weights
<sup>d</sup>using pre-trained GSnet model with optimized GNN weights
<sup>e</sup>using pre-trained GSnet model fine-tuned for residue-level SASA with optimized GNN weights
<sup>f</sup>trained from initialized weights (*i.e.*, not pre-trained).

While the average performance of the doubly fine-tuned GSnet variant (RMSE = 1.29) exceeded the performance of the null model (RMSE = 1.44) and PKAI+ (RMSE 1.30), it fell short when compared to other methods like PROPKA (RMSE = 1.12), CpHMD simulations (RMSE = 1.01), and DeepKa (RMSE = 0.97). This analysis demonstrates that transfer learning targeting more complex residue-level properties based on structural embeddings is possible, but the performance achieved so far lags behind other methods that were specifically developed for p*K_a_* predictions.

### Local charge-aware embedding for pK_a_ predictions

Although GSnet was trained to reproduce geometric, hydrodynamic, and electrostatic solvation free energies, it does not explicitly consider information about local charge distributions which may be essential for making accurate p*K_a_* shift predictions. To test whether a charge-aware local structural embedding can improve p*K_a_* predictions, we developed aLCnet, an atomic variant of our GNN model that is more lightweight than GSnet and tailored specifically to p*K_a_* predictions by including partial atomic charges as an input feature. This model was trained directly on the PHMD549 data to make p*K_a_* predictions and achieved significantly better performance than the GSnet-based models (**Table 2**). The average RMSE of aLCnet on the EXP67S set was 1.10, with the best model achieving an RMSE of 1.08, exceeding the performance of PROPKA (RMSE = 1.12). However, aLCnet still could not reach the reported accuracy of DeepKa (RMSE = 0.97) or CpHMD simulations (RMSE = 1.01). The performance of GSnet and aLCnet can be further compared by looking at the distribution of individual data points (**Fig. S17**). GSnet-based predictions are generally more conservative, closer to the null model, with only few predictions giving shifts of more than 2 p*K_a_* units, and the slope of predicted vs. experimental p*K_a_* values is only about 0.3. With aLCnet, larger shifts are predicted, giving an improved slope of 0.48, similar to DeepKa, but with larger deviations from the experimental values. The significant improvement with aLCnet over GSnet was likely due to the inclusion of atomic charge as an input feature, as well as its truly atomic resolution, highlighting the limitations of using a generic structure-based embedding, such as GSnet trained on global molecular properties, for predictions requiring higher resolution and additional features.

### Accurate predictions of experimental pK_a_ values using embeddings

Training based on calculated p*K_a_* shifts in the section above resulted in good accuracy for predicting experimental p*K_a_* values, especially with aLCnet, but the lack of experimental data in the training set likely limited the performance. On the other hand, training a machine learning model only on experimental p*K_a_* data is problematic because there are only relatively few data points available, especially when eliminating redundant data for the same residue in the same or similar protein structures^43, 44^. Attempting to overcome these challenges, we pursued two distinct transfer learning strategies with GSnet (see **Fig. S6**) and aLCnet (see **Fig. S7**) embeddings that were fine-tuned along with the output MLP by training on experimental p*K_a_* shifts.

In order to train using experimental data, we constructed the MSU-pKa-training, MSU-pKa-validation, and MSU-pKa-test datasets, primarily from entries in PKAD-1^43^ and PKAD-2^44^ as described in the Methods section. The separation between training, validation, and test data was done carefully to avoid redundancy and to prevent data leakage from training to test performance. We note that overlap between training and test data likely resulted in overly optimistic performance estimates in other studies^35^.

Following the transfer learning strategies, we achieved an average RMSE of 1.01 p*K_a_* units with GSnet and an average of 0.95 using aLCnet embeddings, better than values obtained with the null model^66^ (1.21), slightly better than values from the machine learning methods DeepKa^65^ (1.03), PKAI+^67^ (1.01), pKALM^47^ (1.07), better than the physics-based predictor PypKa^68^ (1.03) and only slightly worse than PROPKA^69^ (0.91) (**Table 3**). On the test set, predictions with aLCnet embeddings again achieved a slope of about 0.5 when comparing predicted shifts to experimental shifts (Fig. 5) whereas DeepKa and GSnet-based predictions had shallower distributions due to underpredicting larger shifts (Fig. 5). We note that DeepKa does not predict p*K_a_* shifts for tyrosine and cysteine residues, so performance was evaluated both by applying the null model instead and by excluding tyrosine and cysteine residues (**Table 3**). A direct comparison with other machine learning methods^35, 70^ was not done because of significant overlap between the training sets used in the other methods and the MSU-pKa-test or because code was not available as for the recently published predictors KaML-CBtree and KaML-GAT^46^. However, a similar RMSE value of 1.00 as with our model was reported with the P-SPOC model when testing on sequences that were not included during training^70^. Using pre-trained models resulted in significantly better performance than training from random initial weights. For the GSnet architecture we achieved an RMSE of 1.35 without pre-training, compared to 1.01 with pre-training. For aLCnet we found an RMSE of 1.16 without pre-training, compared to 0.95 with pre-training. These results highlight the significant advantage of transfer learning when training with sparse data.

**Figure 5.**
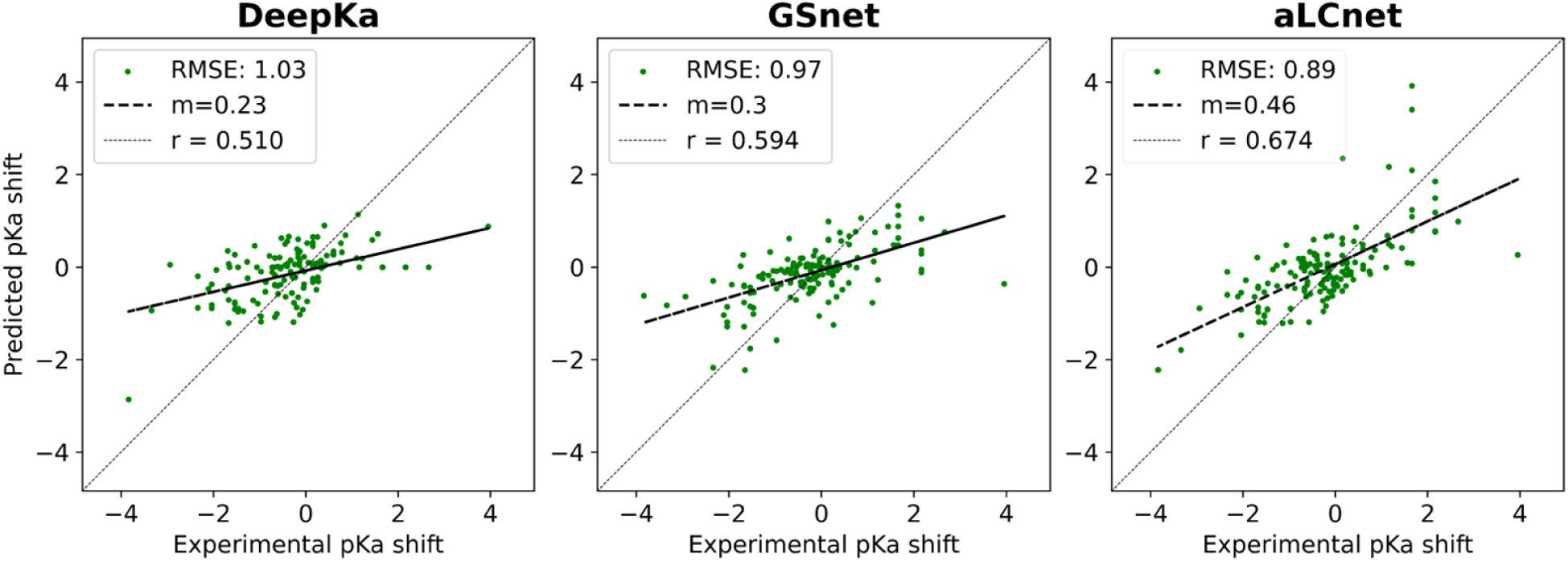
Experimental pKa shift predictions with DeepKa, GSnet, and aLCnet on MSU-pKa-test. For GSnet and aLCnet models the test performance is shown for the model with the lowest validation RMSE (out of 20 trained models). Performance data for DeepKa was obtained via predictions made with the DeepKa web server^65^. DeepKa predicted shifts for tyrosine and cysteine residues were obtained with the null model. RMSE values are given in p*K* units.

**Table 3.** Performance metrics for different models evaluated on MSU-pKa-test. For GearNet, ESM-2, and our models, 20 training runs were conducted and the top 10 models based on validation RMSE were selected for evaluation on the test set; performance results are given as mean ± SEM with the best (out of 10) result given in parentheses. “random” indicates that a model was trained from random initial weights. Predictions with DeepKa were obtained via the DeepKa web server^65^. Predictions with PypKa^68^, PKAI+^67^, and pKALM^47^ were done with software downloaded from the respective github archives described in the publications.

| Method | RMSE | R | m |
| --- | --- | --- | --- |
| Null Model | 1.21 | - | 0.0 |
| PROPKA | 0.91 | 0.57 | 0.69 |
| PypKa | 1.03 | 0.59 | 0.58 |
| DeepKa<sup>a</sup> | 1.03 (0.88 w/o Y/C) | 0.51 (0.59 w/o Y/C) | 0.23 (0.33 w/o Y/C) |
| PKAI+ | 1.01 | 0.54 | 0.27 |
| pKALM | 1.07 | 0.53 | 0.29 |
|  | 1.25<sup>b</sup> | 0.47 | 0.21 |
| GearNet<sup>c</sup> | 1.35 ± 0.01 (1.33) | 0.42 ± 0.01 (0.45) | 0.22 ± 0.00 (0.24) |
| ESM-2<sup>c</sup> | 1.29 ± 0.01 (1.23) | 0.22 ± 0.00 (0.25) | 0.15 ± 0.00 (0.18) |
| GSnet (rSASA-opt)<sup>d</sup> | 1.01 ± 0.00 (0.97) | 0.56 ± 0.00 (0.60) | 0.28 ± 0.00 (0.38) |
| GSnet (random) | 1.35 ± 0.00 (1.28) | 0.43 ± 0.01 (0.52) | 0.19 ± 0.00 (0.23) |
| aLCnet (opt)<sup>e</sup> | 0.95 ± 0.00 (0.89) | 0.62 ± 0.00 (0.68) | 0.38 ± 0.01 (0.59) |
| aLCnet (random) | 1.16 ± 0.01 (1.02) | 0.39 ± 0.02 (0.56) | 0.04 ± 0.01 (0.19) |

<sup>a</sup>Predictions for tyrosine and cysteine residues obtained via null model, metrics excluding these residues shown in parentheses.
<sup>b</sup>For a subset of 102 out of 143 data points without redundancy to pKALM training set<sup>47</sup>
<sup>c</sup>Embeddings passed to the same MLP architecture as in our models.
<sup>d</sup>Pre-trained with six original physicochemical properties + residue-level SASA and optimized
<sup>e</sup>Pre-trained with CpHMD pK<sub>a</sub> data from PHMD549 dataset<sup>36</sup> and optimized

We also evaluated performance of the models by residue type, and these results are provided in **Table S5**. For aspartate residues, PROPKA and aLCnet performed similarly well with RMSEs near 0.9, with GSnet performing worse than the null model. All models, including the null model, performed similarly well for glutamate, with an RMSE around 0.6, except aLCnet with an RMSE of about 0.7. GSnet performed exceptionally well for histidine (RMSE = 0.73), with PROPKA (RMSE = 1.22) and aLCnet (1.15) performing worse. For lysine residues, aLCnet, PROPKA, and the null model achieved RMSEs near 0.5, while GSnet achieved RMSEs near 0.95. GSnet and aLCnet outperformed the null model (RMSE = 1.72) for tyrosine residues with RMSEs of 1.11 and 1.08, respectively, but they were slightly worse than PROPKA (RMSE = 0.82). GSnet performed exceptionally well for cysteine residues with an RMSE of 0.11, and aLCnet was able to exceed the performance of the null model (RMSE = 1.65) as well with an RMSE of 1.03. PROPKA performed quite poorly for cysteine with an RMSE of 4.10.

We tested whether alternative pre-trained embedding models, namely GearNet and ESM-2, could perform similarly well as GSnet and aLCnet. With GearNet we achieved an average RMSE of 1.35 while ESM-2 embeddings appeared to work slightly better, with an average RMSE of 1.29. We note that our results using ESM-2 embeddings are consistent with the performance of pKALM^47^, which also uses ESM-2 embeddings, when applied to the subset of our test cases that are non-redundant to the training set of pKALM^47^ according to our similarity classification strategy. We speculate that that the significantly better test performance reported in the pKALM paper may be due to data leakage between training and test sets. Based on our results, it seems that neither GearNet nor ESM-2 embeddings allow for accurate p*K_a_* predictions, as neither model, even when selecting the best of all trained models, was able to exceed the performance of the null model (RMSE = 1.21).

### Predictions of experimental pK_a_ values in IDPs

We further tested the performance of GSnet and aLCnet-based p*K_a_* predictions values by applying these models to α-synuclein, an IDP. Predictions were compared with experimental p*K_a_* measurements for 18 glutamate residues, 6 aspartate residues, and 1 histidine residue^71^. Because the structure of an IDP is inadequately described by a single structure^72^, we made predictions over an ensemble of 300 simulated structures and averaged the predictions for each residue. We found that the averaged p*K_a_* value predictions were generally in good agreement with the experimental values, with RMSE values of 0.20 and 0.45 for GSnet and aLCnet predictors, respectively, across all 25 residues for which experimental p*K_a_* values are available (Fig. 6). The slightly worse performance with aLCnet was due to systematically underestimating most of the p*K_a_* values compared to the experimental values, whereas GSnet did not show such a trend. This could indicate that using local embeddings based on folded proteins may not transfer as well to residues in IDPs as global structure embeddings. Additionally, it is possible that the better performance of GSnet could be attributed to its pre-training on a more diverse dataset of protein structures, which likely included a broader range of disordered and flexible regions.

**Figure 6.**
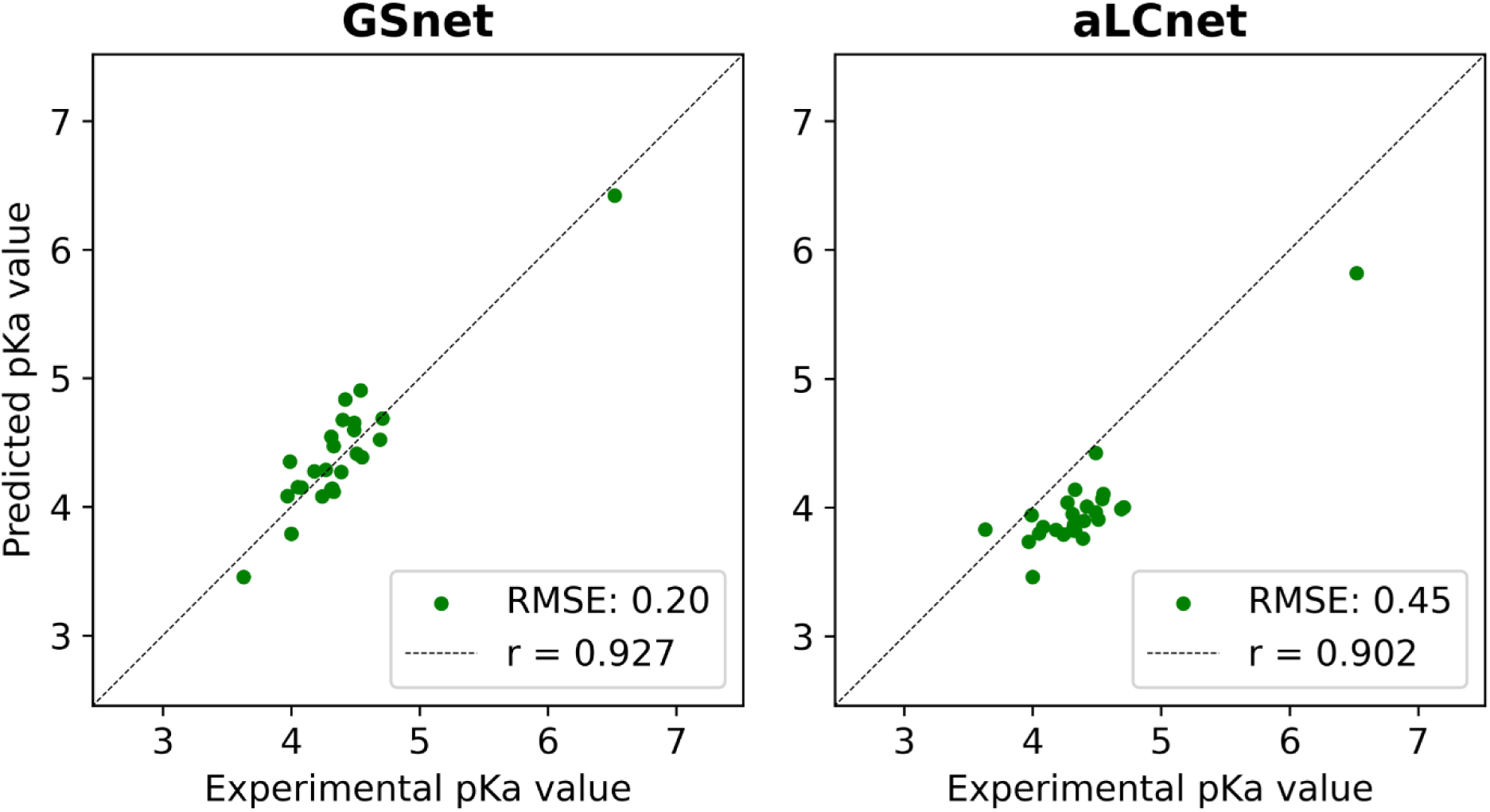
GSnet and aLCnet predictions of p*K_a_* values for selected residues in α-synuclein. Experimental values were obtained via NMR in 150 mM NaCl by Croke *et al.*^71^. RMSE values are given in p*K* units.

### High-throughput Predictions

Once trained, machine learning frameworks are very fast, especially when making multiple predictions, as they can be made in parallel on GPU computing hardware. Thus, a significant advantage of GSnet, and the related aLCnet, is vastly reduced computational cost. For instance, when making global property predictions for 4-aminobutyrate aminotransferase (UNIPROT ID P80404), a 500-residue protein, it takes more than 4 minutes to produce results with HYDROPRO and more than 45 minutes to obtain solvation free energies with APBS, whereas the trained GSnet model only requires about 1 second to make a forward pass. (**Table S6**). Apart from faster predictions for 4-aminobutyrate aminotransferase, GSnet also uses much less memory, about 670 MB, compared to 1.7 GB with HYDROPRO and 30.8 GB with APBS (**Table S6**). The fast calculation of electrostatic solvation free energies is especially interesting in the application of continuum solvation models, whose applications have been plagued by the relatively high cost of obtaining solvation free energies using traditional approaches^73^. However, further efforts are needed to increase the accuracy of solvation free energy predictions for practical applications, which is beyond the scope of the present work.

When considering p*K_a_* predictions, it took about 180 seconds with GSnet and 130 seconds with aLCnet to predict 2000 residues on one NVIDIA GeForce RTX 2080 Ti GPU, and the cost with aLCnet includes the time to perform hydrogen bond optimization and to generate input charges via PDB2PQR, which can likely be optimized in a future version. For comparison, predictions with PROPKA take about 231 seconds for 2000 residues. This opens up high-throughput p*K_a_* predictions for very large complexes or for a large number of structures. To illustrate a possible application, we calculated p*K_a_* shifts for all ionizable residues in the GroEL-GroES chaperonin complex (Fig. 7). The resulting shifts mapped onto the complex structure show distinct patterns of p*K_a_* shifts, for example significant shifts of basic residues near the conical top towards acidic pH values (Fig. 7).

**Figure 7.**
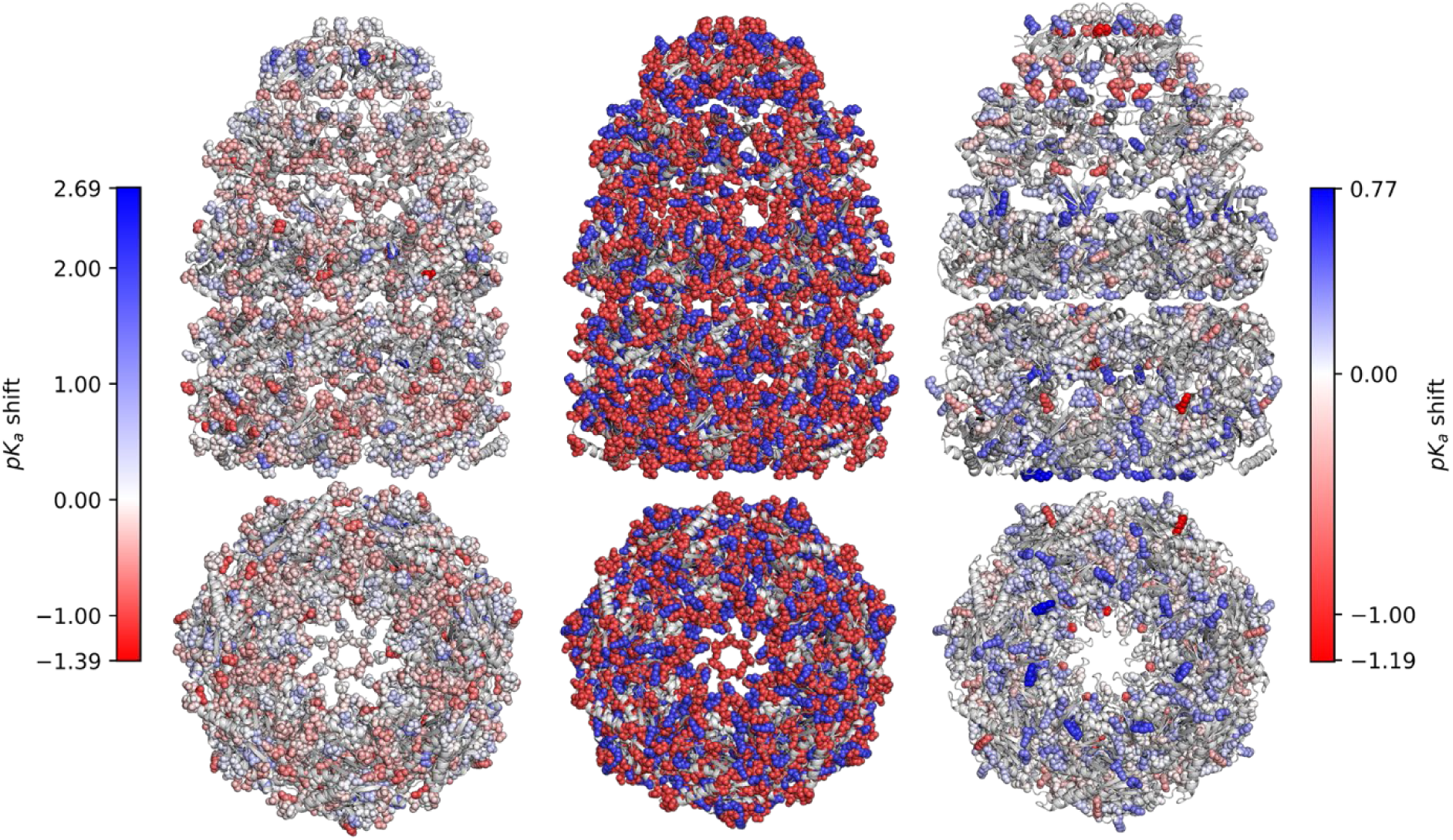
Predicted p*K_a_* shifts for GroEL-GroES by aLCnet. Predicted shifts of acidic and basic residues are shown on the left and right, respectively. The color scale indicates the magnitude of the shift, with deeper shades of blue indicating a greater increase in p*K_a_*, deeper shades of red indicating a greater decrease in p*K_a_*, and white indicating no shift. The central panel indicates all ionizable residues, with acidic residues colored red and basic residues colored blue. The figure was generated using PyMol^74^.

## CONCLUSIONS

The work presented here demonstrates the utility of GNNs like GSnet and aLCnet in predicting physicochemical properties of proteins while simultaneously generating molecular embeddings that can be employed in the prediction of other properties via transfer learning. We show that using embeddings and transfer learning results in better model performance, especially when training on new properties for which only sparse data is available. We demonstrate successful transfer learning to predict solvent-accessible surface areas, and we make p*K_a_* shift predictions at accuracies that are competitive with methods developed specifically for such applications. Moreover, using GSnet and aLCnet embeddings, we achieve better accuracy than using previously developed embeddings, namely ESM-2 and GearNet.

We find good performance using GSnet, a global structure embedding, but the best performance for p*K_a_* predictions required the higher resolution, charge-aware aLCnet. In principle, it should be possible to learn atomistic features together with global structural properties, and future work may focus on generating a more comprehensive embedding that better bridges different scales. Presumably, such an embedding would involve a higher-capacity network and would need to be trained on a larger variety of training data. For example, one could imagine training such an embedding on local electrostatic properties or properties that capture local solvation environments such as Poisson Boltzmann-dervied Generalized Born radii^75^. Additionally, future work may benefit from incorporating conformational dynamics, as experimental properties constitute ensemble averages over thermally fluctuating molecules, rather than the static structures considered here.

As for the accurate prediction of p*K_a_* values, we find very good performance based on aLCnet embeddings that matches the accuracy of other empirical methods but may not yet surpass the accuracy of simulation-based constant-pH approaches^10, 76^. The main limitation is likely the limited amount of experimental data available for training, especially when considering the need for non-redundant data for developing transferable models that perform well beyond the training set. Using computational data for training appears to partially resolve the issue as demonstrated previously^36, 65^, but to make further progress with ML-based p*K_a_* predictors, much more extensive experimental data is probably needed.

On the practical level, our aLCnet-based p*K_a_* predictor is computationally very efficient, offering accuracy similar to PROPKA but at greater speed. This opens up new applications where p*K_a_* shift predictions could be integrated into structural analysis pipelines. It is also becoming possible now to apply p*K_a_* shift predictions to very large complexes and to large numbers of structures up to proteome-wide analyses.

## ACKNOWLEDGEMENTS

We thank Lim Heo for his valuable insights regarding ESM-2 and help in generating the IDP data. We also acknowledge discussions with Alexander Jussupow and Gilberto Valdes-Garcia. Funding was provided by the National Institute of Health (NIGMS) grant R35 GM126948 and by the National Science Foundation, grant MCB 2210228.

## SUPPORTING MATERIAL

Supporting Tables S1-S6, Figures S1-S17, and supplementary references related to those tables and figures are provided as supporting material. This information is available free of charge via the Internet at http://pubs.acs.org

## TOC GRAPHIC

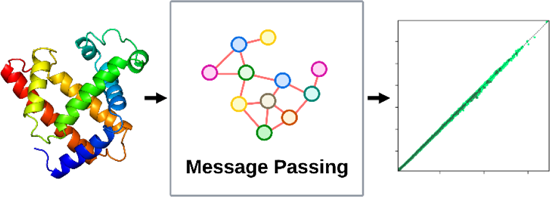

