## Supplementary Material for "Accurate Predictions of Molecular Properties of Proteins via Graph Neural Networks and Transfer Learning"

for

**Figures S1-S17**
**Tables S1-S6**
**Supplementary References**

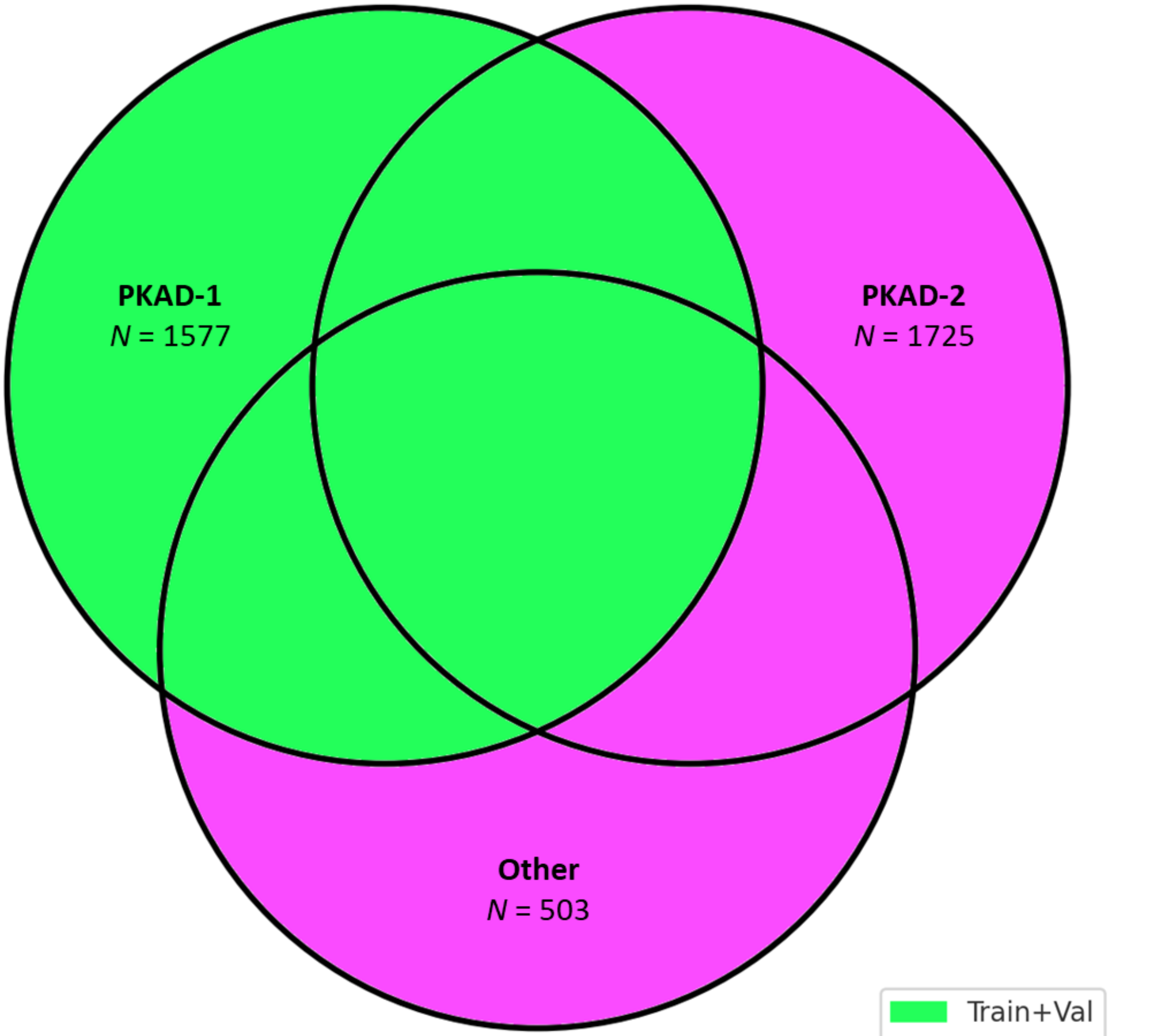


**Figure S1.** Visual depiction of the sources of our experimental datasets. "Other" includes experimental pKa data that was obtained from the experimental references listed in Chen *et al*.[1] Gokcan *et al*.[2], and Wilson *et al*.[3] that were not obtained directly from PKAD. Overlap in the Venn diagram represents data points that were considered similar according to the similarity classification strategy outlined in the Methods section. Data points outside of PKAD-1 were only added to the test set if they were dissimilar to data points in the training/validation set according to the similarity classification strategy. *N* represents the total number of data points in each data source.

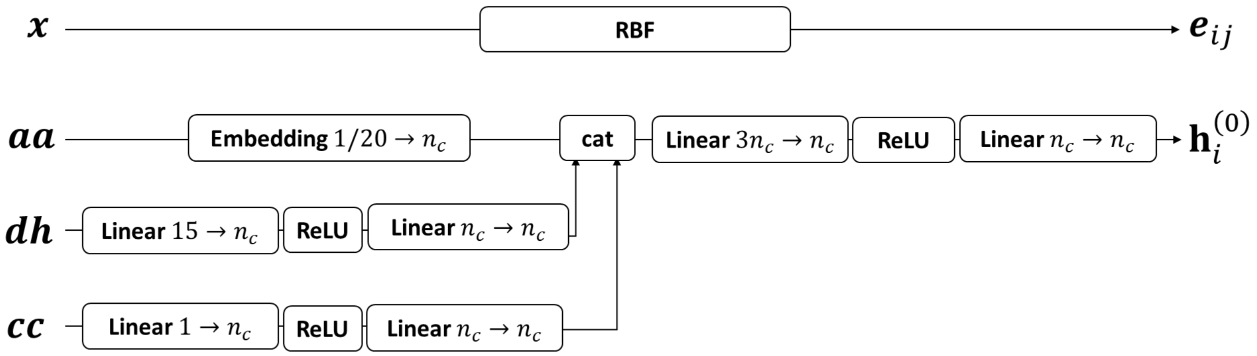


**Figure S2.** GSnet node and edge embeddings from input features. ($n_c$: number of hidden channels, RBF: radial basis function, cat: concatenate)

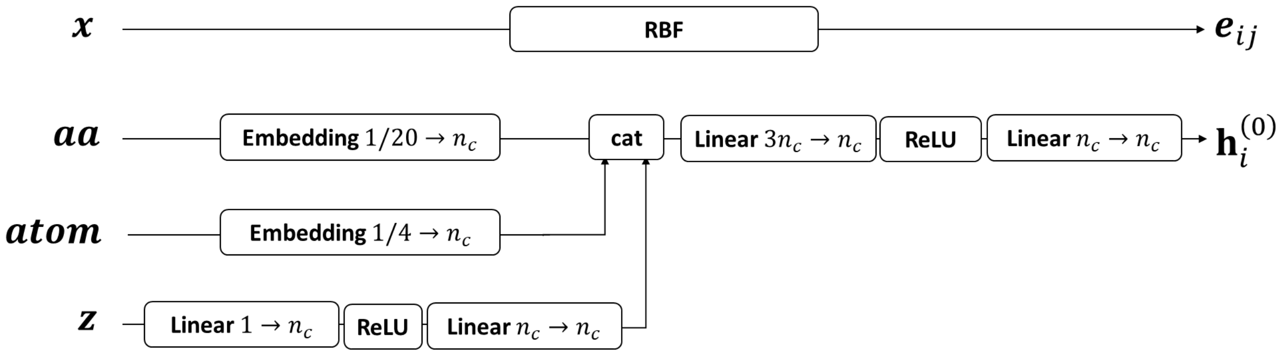


**Figure S3.** aLCnet node and edge embeddings from input features. ($n_c$: number of hidden channels, RBF: radial basis function, cat: concatenate, $z$: charge)

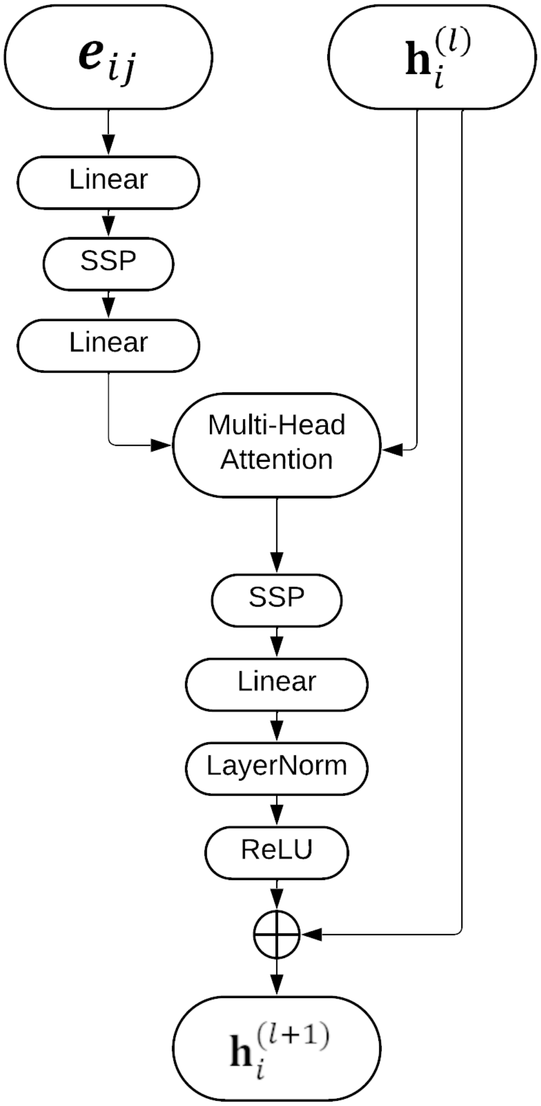


**Figure S4.** Architecture of message passing blocks in GSnet and aLCnet. $\mathbf{h}_i^{(l)}$ are node features of node $i$ at layer $l$, $\boldsymbol{e}_{ij}$ are edge features between neighboring nodes $i$ and $j$, and $\mathbf{h}_i^{(l+1)}$ are node features of node $i$ at the next layer $l+1$. SSP is a ShiftedSoftPlus activation function defined as $\mathrm{SSP}(\mathbf{x}) = ln(\mathbf{1} + e^{\mathbf{x}}) - ln(2)\mathbf{1}$. The operations of the Multi-Head Attention mechanism are described by Shi *et al.*[4]. LayerNorm refers to the layer normalization operation described by Lei Ba *et al.*[5].

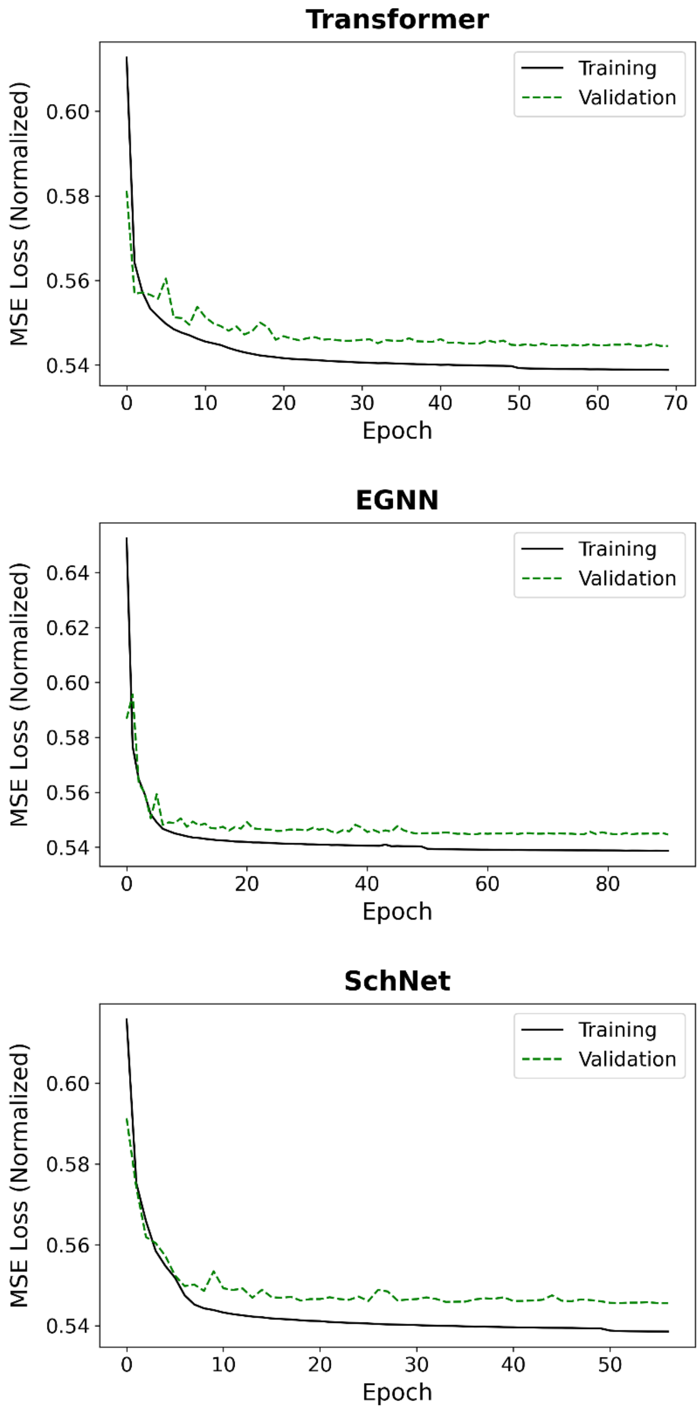


**Figure S5.** Mean-squared error (MSE) loss across the six target values as a function of epoch during training. Training loss is shown as a solid black line and validation loss is shown as a dashed green line.

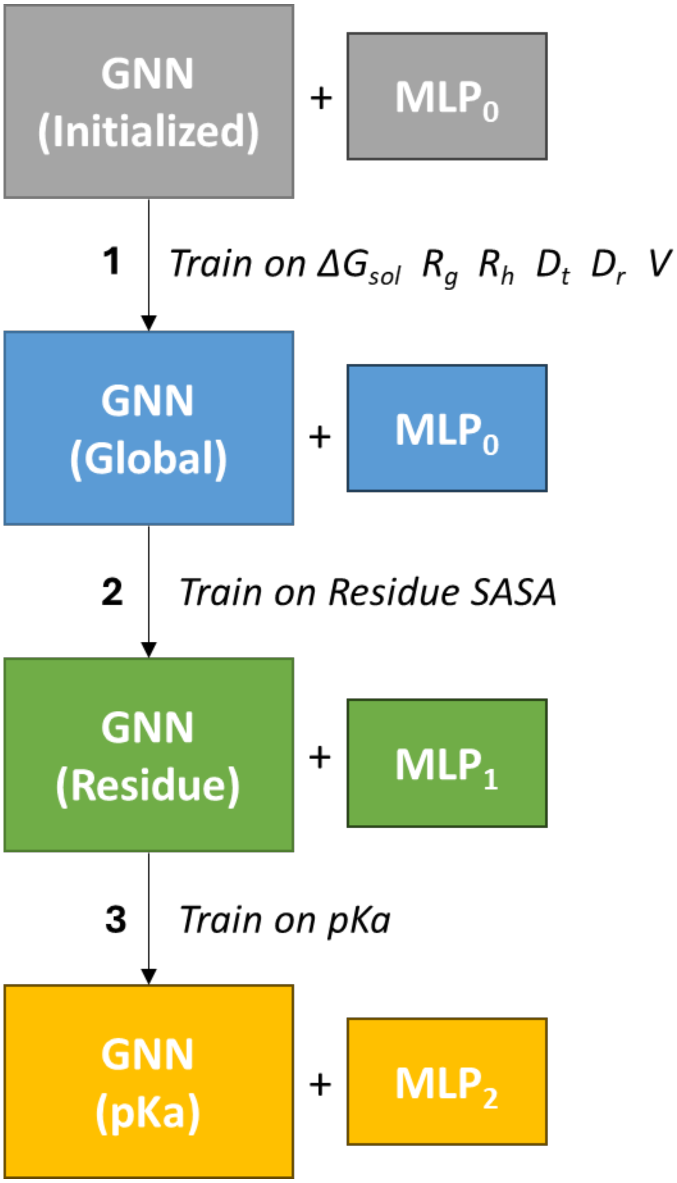


**Figure S6. GSnet training scheme for p$K_a$ transfer learning.** First, the initialized GNN and MLP were trained on the original target values. Next, this pretrained GNN and a new MLP were further trained to make $SASA_{res}$ predictions. Finally, the GNN as pretrained for $SASA_{res}$ was transferred, and yet another new MLP was introduced and trained to make p$K_a$ predictions. Note that the same process was used for training on both p$K_a$ datasets.

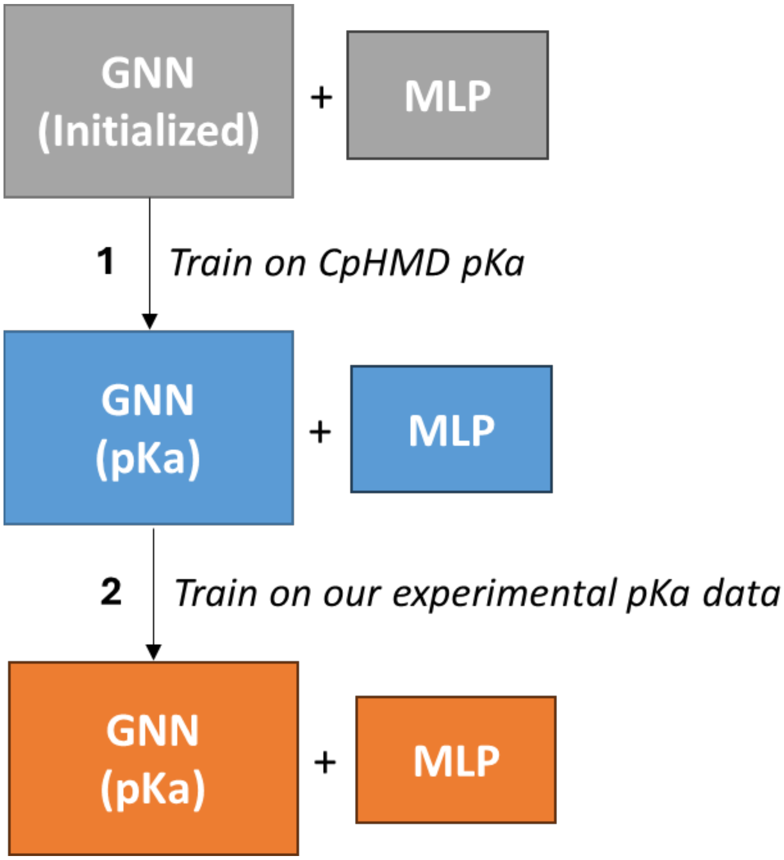


**Figure S7. aLCnet training scheme for p$K_a$ transfer learning.** First, the initialized GNN and MLP were trained on the simulated p$K_a$ data in the PHMD547 dataset. Next, this pretrained GNN and MLP were further trained on our experimental dataset.

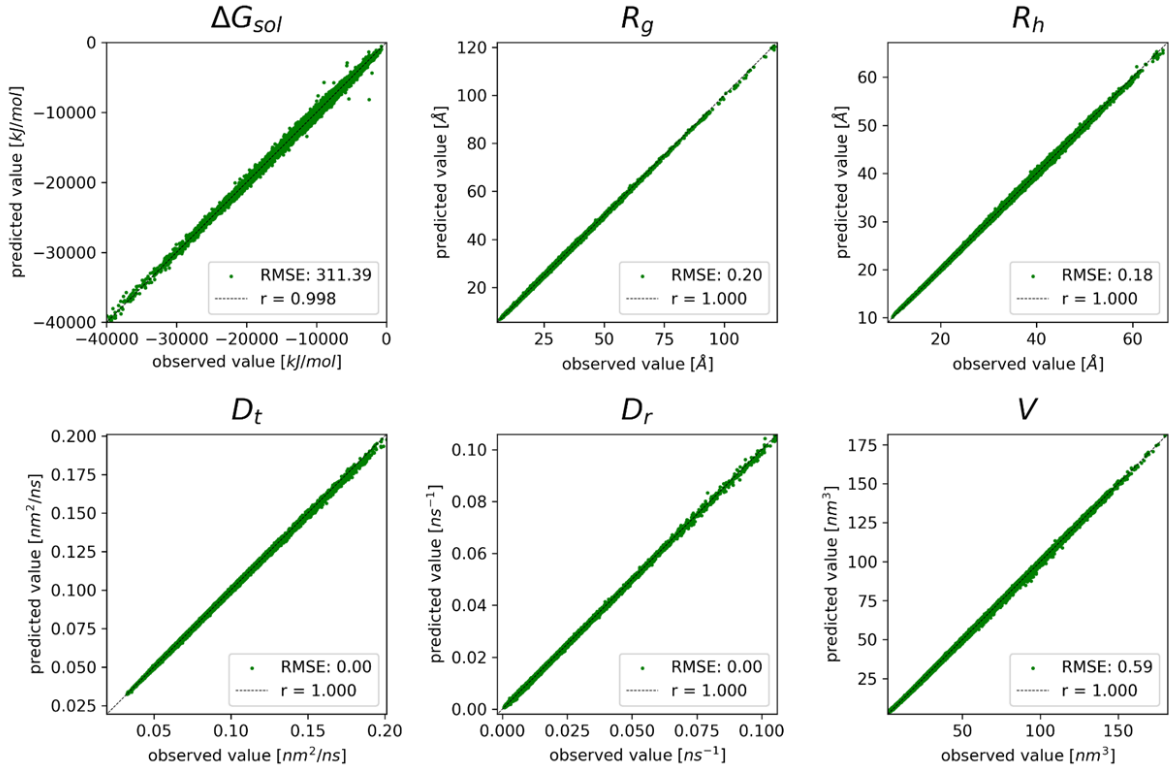


**Figure S8. Training set performance of transformer model for global molecular properties vs. reference values**. Results are shown for $\Delta G_{sol}$, $R_g$, $R_h$, $D_t$, $D_r$, and $V$. RMSE values are given in units as displayed on the axes of each subplot.

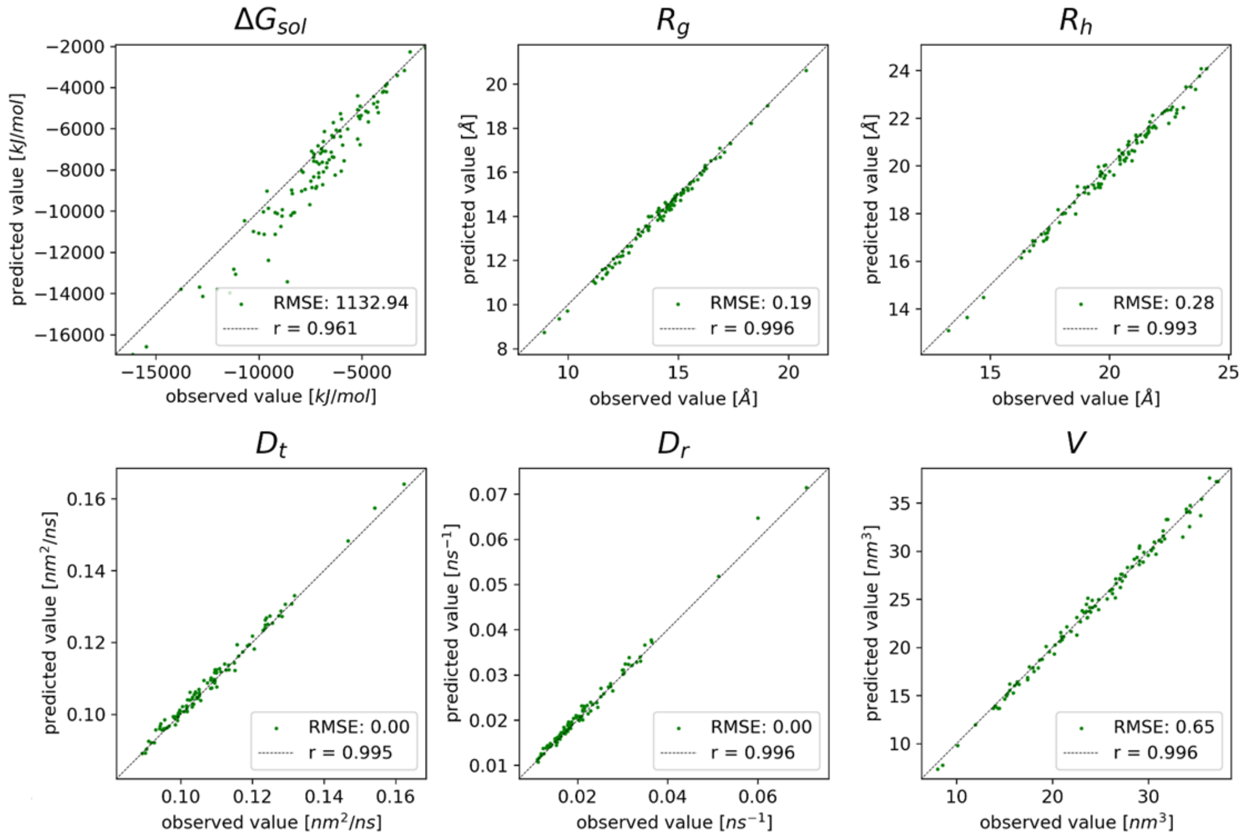


**Figure S9. GSnet performance for predictions of global molecular properties on small protein test set.** Predictions are shown for 123 dissimilar (<20% identity), small (40-200 residues) PDB structures. (chosen via PISCES[6]; resolution < 1.5 Å, R<0.25). RMSE values are given in units as displayed on the axes of each subplot.

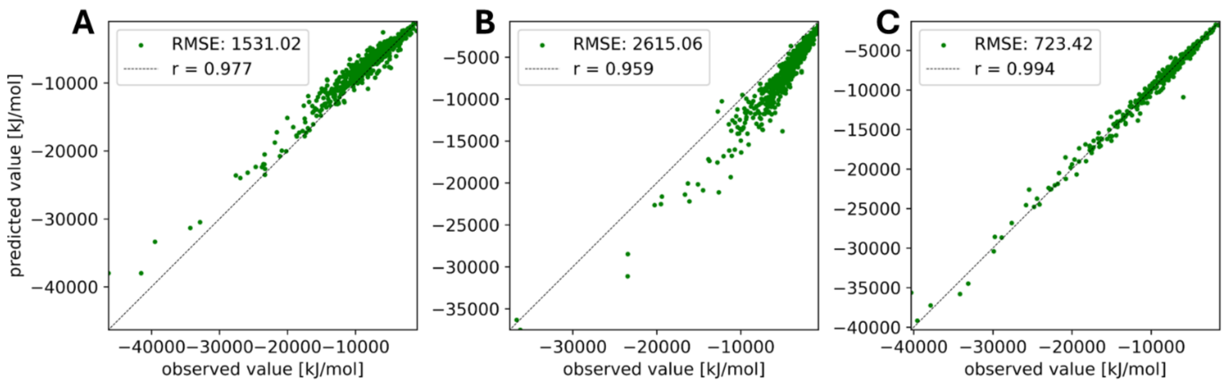


**Figure S10. GSnet performance for predictions of $\Delta G_{sol}$ on extended PDB test set.** Predictions are shown for 610 proteins without (A) and with (B) energy minimization. Results with AlphaFold2[7] models (C) are shown for comparison. RMSE values are given in kJ/mol.

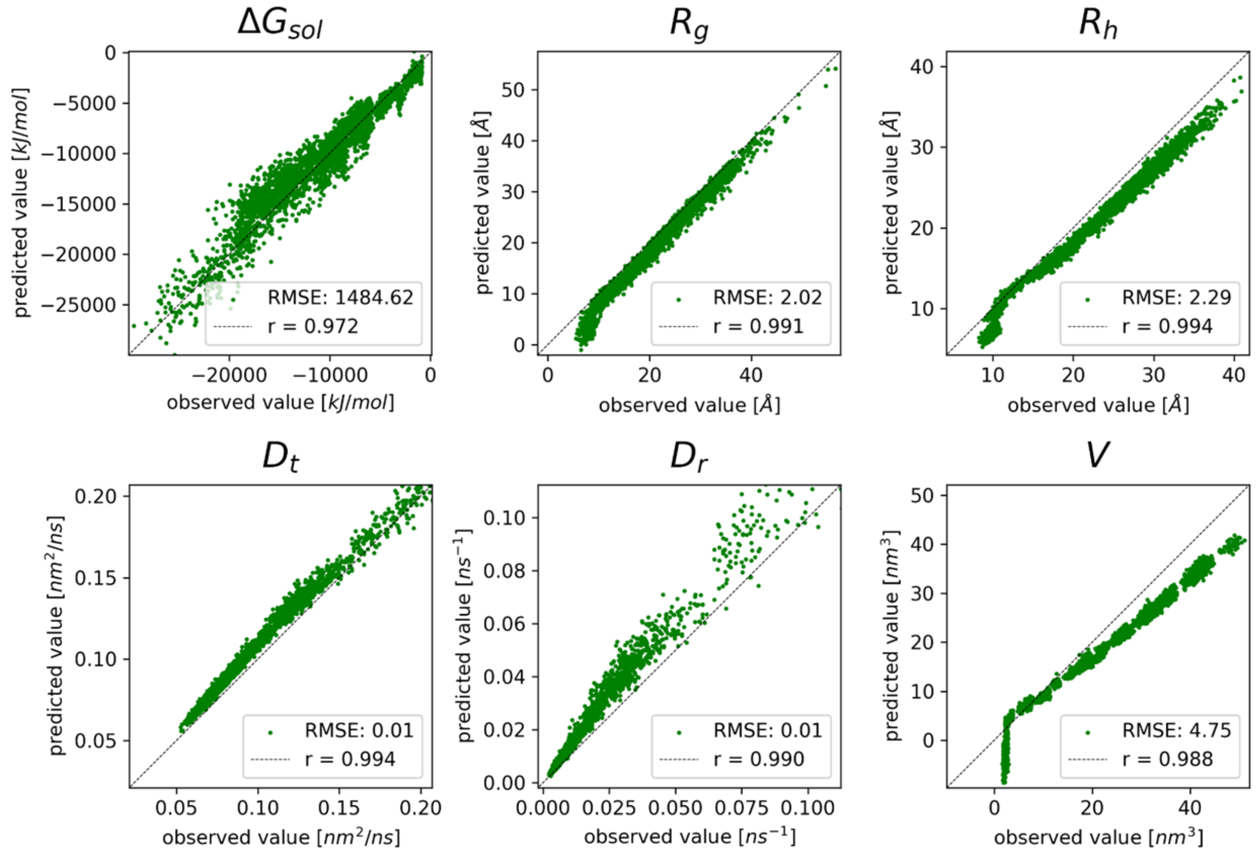


**Figure S11. GSnet performance on IDP structures.** This validation set consists of 4,500 total IDP structures (ensembles of 100 structures for 45 distinct peptides; generated via *cg2aa*). RMSE values are given in units as displayed on the axes of each subplot.

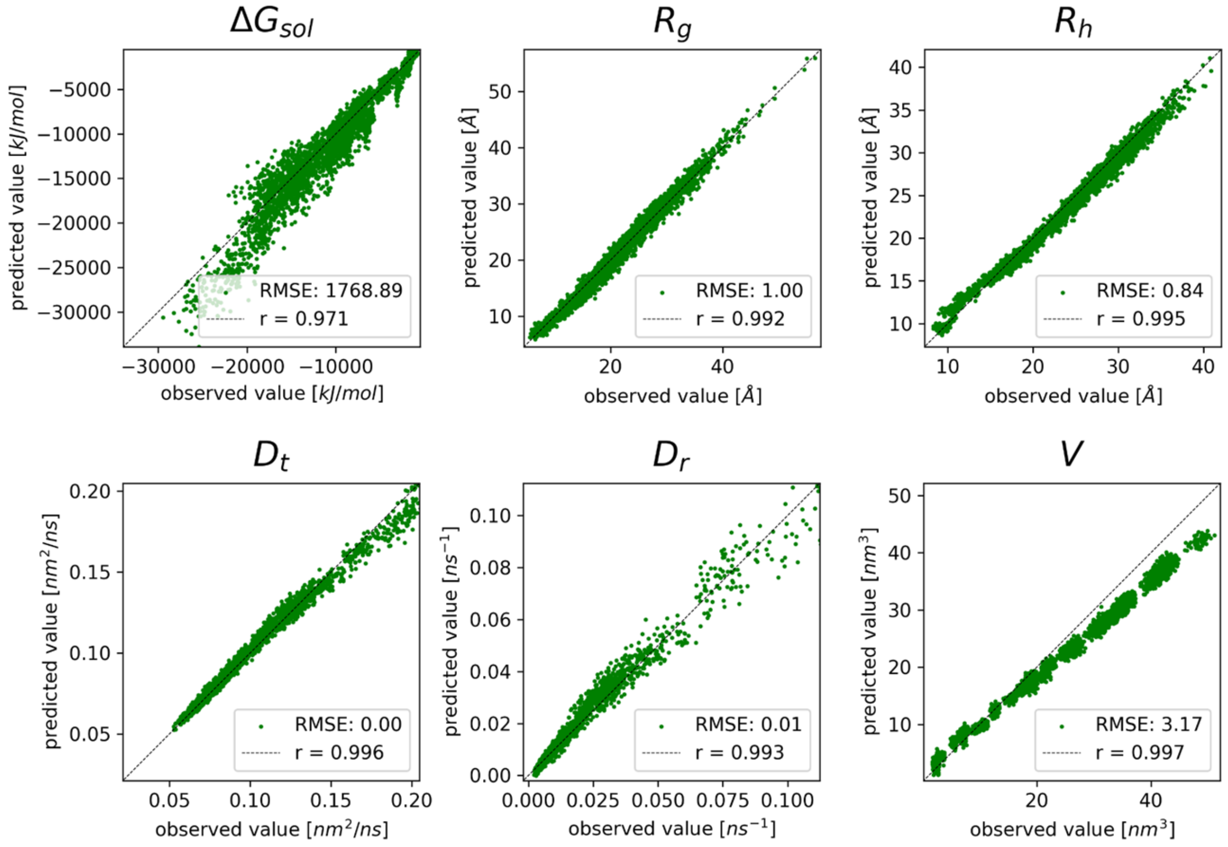


**Figure S12. Retrained GSnet performance on IDP structures.** Performance of GSnet after fine-tuning on a training set consisting of 100 angiotensin structures. The validation set consists of 4,500 total IDP structures (ensembles of 100 structures for 45 distinct peptides; generated via *cg2aa*). RMSE values are given in units as displayed on the axes of each subplot.

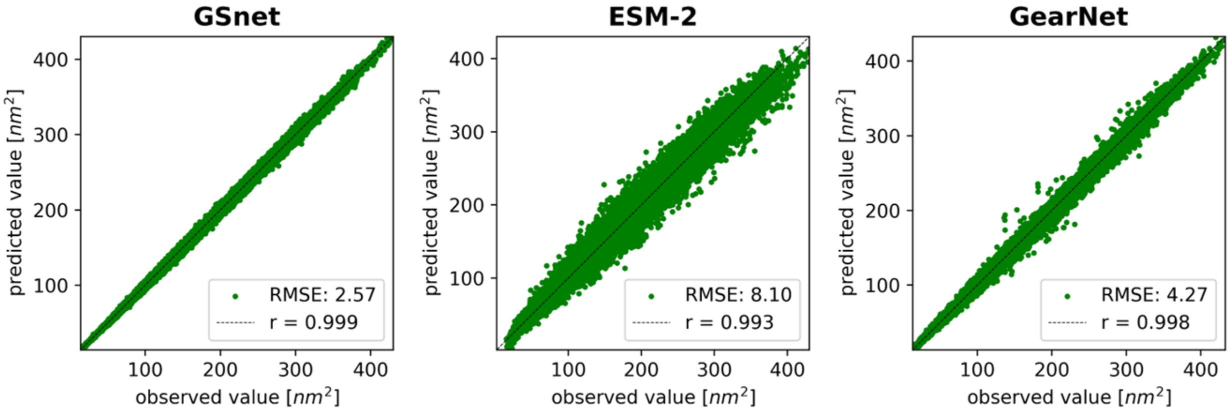


**Figure S13. Training set performance of embedding-based SASA predictors.** Correlation coefficients from linear fits are shown in the upper left corner of each plot. RMSE values are given in $nm^2$.

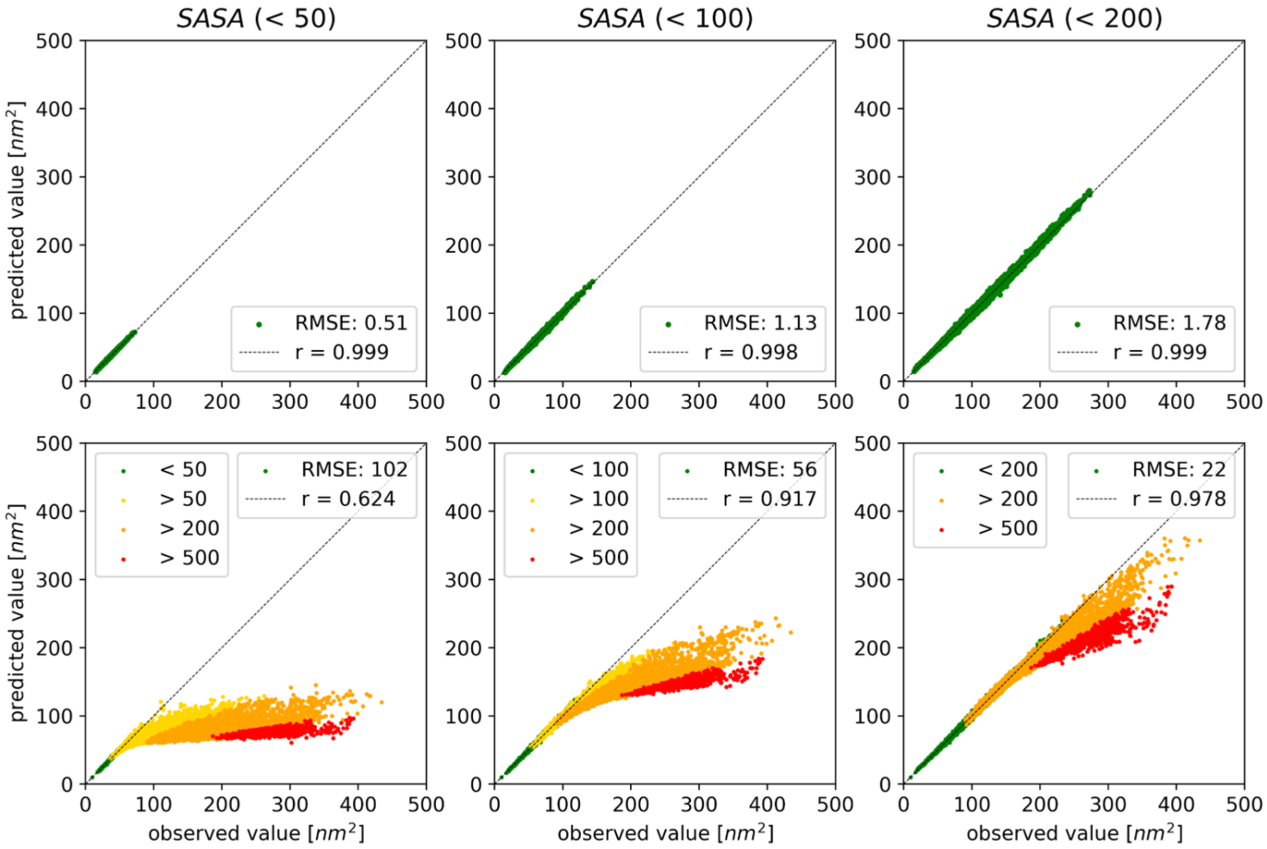


**Figure S14. GSnet-based SASA predictors trained on proteins with limited size.** Model performance is shown for training (top row) and validation (bottom row) sets. Numbers in parentheses indicate maximum size (in residues) of structures in training set. Each data point represents one protein, with colors signifying the protein size as indicated in the figure legend. Pearson correlation coefficients and RMSEs encompass all data shown in the plots. RMSE values are given in $nm^2$.

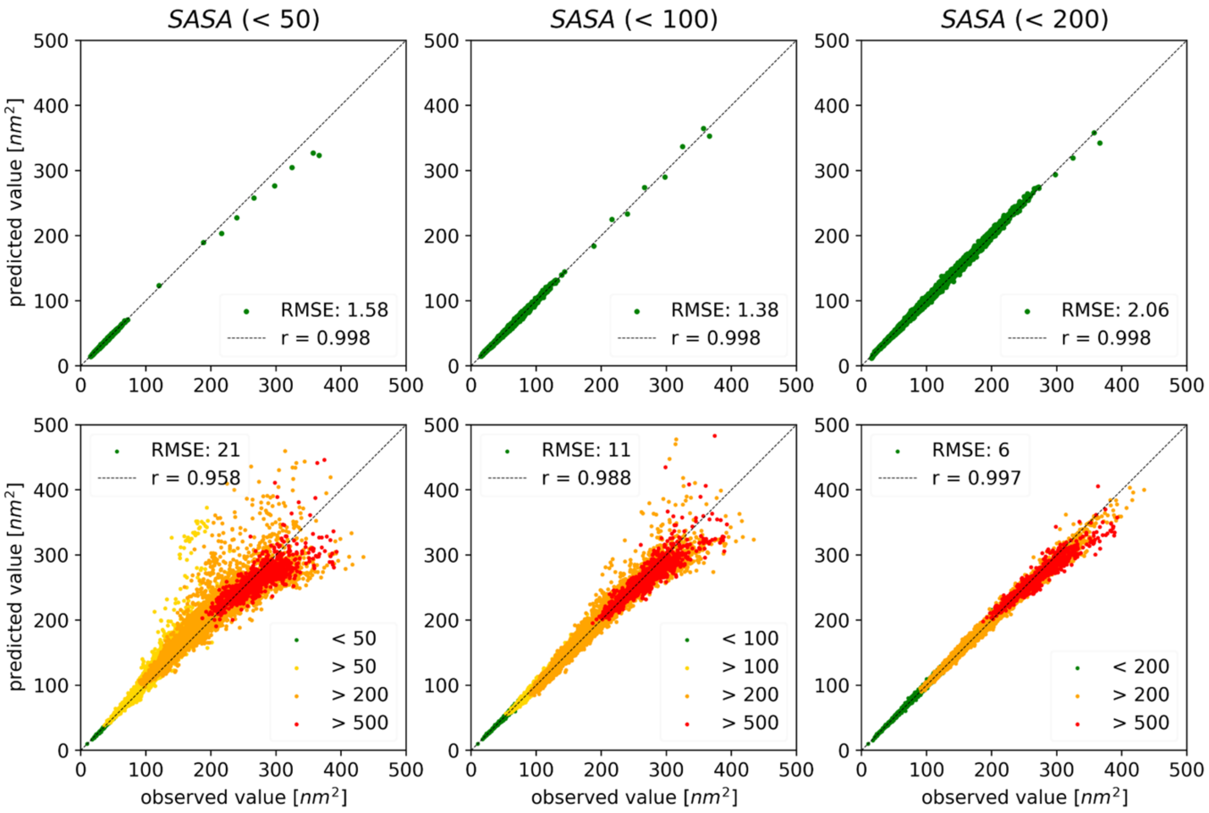


**Figure S15. GSnet-based SASA predictors trained on proteins with limited size and augmented data for larger proteins.** Model performance is shown for training (a) and validation (b) sets. Numbers in parentheses indicate maximum size (in residues) of structures in training set, excluding augmented data points. Each data point represents one protein, with colors signifying the protein size as indicated in the figure legend. Pearson correlation coefficients and RMSEs encompass all data shown in the plots. RMSE values are given in $nm^2$.

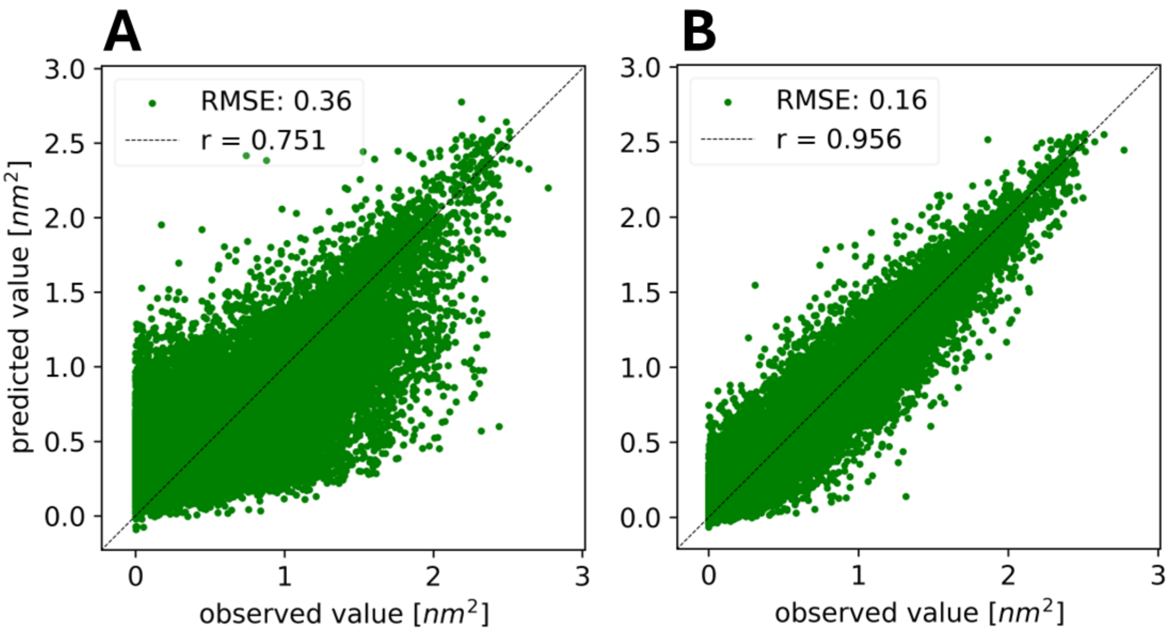


**Figure S16. GSnet-based residue-level SASA predictions.** Plots display model predictions on the validation set vs. calculated values when using: (A) a fixed, pretrained GNN where only the output MLP was trained, and (B) a pretrained GNN where both the GNN parameters and the output MLP were optimized during training. Correlation coefficients from linear fits and RMSE values are shown in the lower right corner of each plot. RMSE values are given in $nm^2$.

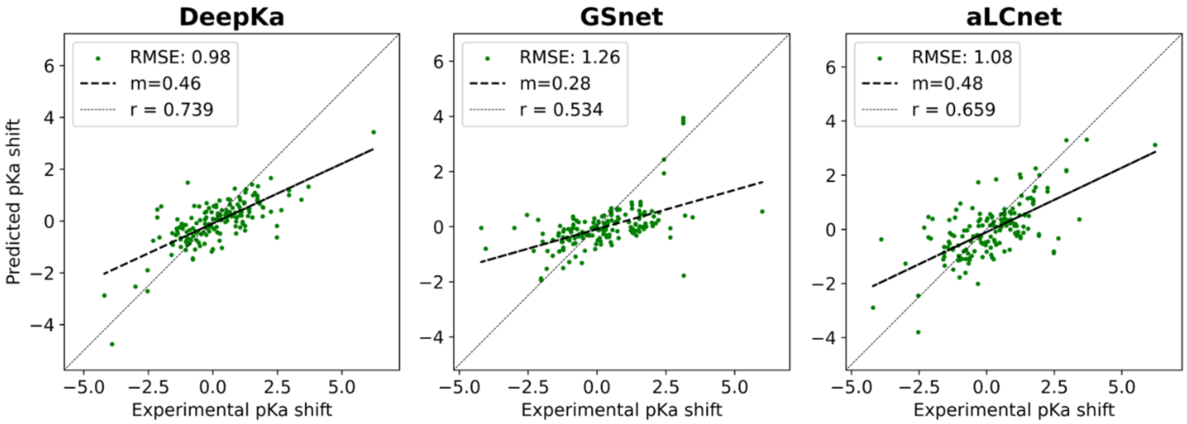


**Figure S17. Experimental pKa shift predictions for EXP67S test set with DeepKa, GSnet, and aLCnet.** For GSnet and aLCnet models the test performance is shown for the model with the lowest validation RMSE (out of 100 trained models). Performance data for DeepKa was obtained via predictions made with the DeepKa Web Server[8]. RMSE values are given in p*K* units.

**Table S1. Input features to GSnet.**

| **Feature** [Shape] | **Description** |
|---|---|
| **x** $[N, 3]$ | Cartesian coordinates of $C_\alpha$ atoms. |
| **aa** $[N, 20]$ | Integer encoding of amino acid sequence. |
| **dh** $[N, 15]$ | Two backbone dihedral angles ($\varphi$,$\psi$) and the first three side-chain dihedral angles ($\chi_1$, $\chi_2$, $\chi_3$) represented as mask, cosine & sine encoding. |
| **cc** $[N, 1]$ | Distances from $C_\alpha$ atoms to protein center of mass. |

*N* is the number of residues in the input protein

**Table S2. Input features to aLCnet.**

| **Feature**<br>[Shape] | **Description** |
|---|---|
| **x**<br>$[N, 3]$ | Cartesian coordinates of heavy atoms. |
| **aa**<br>$[N, 20]$ | Integer encoding of amino acid type. |
| **atom**<br>$[N, 4]$ | Integer encoding of atom type. |
| **charge**<br>$[N, 1]$ | Effective charge of atom as computed by PDB2PQR. |

$N$ is the number of atoms in the input selection

**Table S3. Graph features used as input to MLP.** Features vary according to the predicted properties and are extracted after message passing. "Pretrain" refers to the original training on $\Delta G_{sol}$, $R_g$, $R_h$, $D_t$, $D_r$, $V$. "SASA" refers to fine-tuning for molecular-level SASA, "rSASA" refers to fine-tuning for residue-level SASA, and "p$K_a$" refers to training on p$K_a$ values. $d$ is the number of hidden dimensions of the GNN, $N$ is the number of features extracted with dimensionality $d$, and $d \cdot N$ is the number of input channels to the MLP.

| **Model** | **Task** | **MLP input Features** | $N$ | $d \cdot N$ |
|---|---|---|---|---|
| **GSnet** $d = 150$ | Pretrain | $\boldsymbol{\mu}$ | 1 | 150 |
| | SASA | $\boldsymbol{\mu}$ | 1 | 150 |
| | rSASA | $\mathbf{h}_i$ | 1 | 150 |
| | p$K_a$ | $\text{concat}(\mathbf{h}_i, \boldsymbol{\mu}_6, \boldsymbol{\mu}_8, \boldsymbol{\mu}_{10}, \boldsymbol{\mu}_{12}, \boldsymbol{\mu}_{15}, \boldsymbol{\mu})$ | 7 | 1,050 |
| **aLCnet** $d = 75$ | p$K_a$ | $\text{concat}(\mathbf{h}_{C\alpha}, \boldsymbol{\mu}_{aa}, \boldsymbol{\mu})$ | 3 | 225 |

**Table S4. Training set performance for prediction of global molecular properties.** Mean absolute percent errors (MAPE) and root mean square errors (RMSE) are reported as the mean and standard errors from 20 independent training runs (except for ESM-2).

| | Model | $\Delta G_{sol}$ [kJ/mol] | $R_g$ [Å] | $R_h$ [Å] | $D_t$ [$nm^2/\mu s$] | $D_r$ [$\mu s^{-1}$] | V [$nm^3$] |
|---|---|---|---|---|---|---|---|
| **MAPE (%)** | *Transformer (GSnet)* | 3.29 ± 0.09 | 0.72 ± 0.02 | 0.53 ± 0.01 | 0.53 ± 0.01 | 2.10 ± 0.04 | 1.04 ± 0.01 |
| | *EGNN* | 2.81 ± 0.07 | 0.76 ± 0.02 | 0.50 ± 0.01 | 0.51 ± 0.01 | 2.05 ± 0.04 | 1.03 ± 0.01 |
| | *SchNet* | 2.03 ± 0.04 | 0.71 ± 0.01 | 0.41 ± 0.01 | 0.41 ± 0.01 | 1.89 ± 0.04 | 1.02 ± 0.01 |
| | *GearNet* | 3.79 ± 0.05 | 1.57 ± 0.01 | 0.98 ± 0.01 | 0.98 ± 0.01 | 3.85 ± 0.04 | 2.50 ± 0.02 |
| | *ESM-2*[a] | 8.64 | 4.60 | 2.65 | 2.65 | 4.60 | 5.12 |
| **RMSE** | *Transformer (GSnet)* | 393.6 ± 11.7 | 0.25 ± 0.01 | 0.21 ± 0.01 | 0.55 ± 0.01 | 0.23 ± 0.01 | 0.61 ± 0.01 |
| | *EGNN* | 336.2 ± 9.1 | 0.25 ± 0.01 | 0.20 ± 0.01 | 0.53 ± 0.01 | 0.22 ± 0.00 | 0.58 ± 0.00 |
| | *SchNet* | 231.7 ± 3.6 | 0.22 ± 0.00 | 0.15 ± 0.00 | 0.43 ± 0.01 | 0.21 ± 0.00 | 0.58 ± 0.01 |
| | *GearNet* | 456.4 ± 4.5 | 0.54 ± 0.01 | 0.39 ± 0.00 | 0.96 ± 0.01 | 0.38 ± 0.01 | 1.67 ± 0.01 |
| | *ESM-2*[a] | 1145.0 | 1.85 | 1.20 | 2.91 | 1.36 | 3.46 |

[a]Only one model was trained with ESM-2 due to computational cost.

**Table S5.** Test RMSE by residue type for the various pKa prediction methods. GearNet, ESM, GSnet and aLCnet models selected based on lowest validation RMSE.

| **Model** | ASP | GLU | HIS | LYS | TYR | CYS |
|---|---|---|---|---|---|---|
| *Null Model* | 1.13 | 0.58 | 1.83 | 0.51 | 1.72 | 1.65 |
| *PROPKA* | 0.91 | 0.64 | 1.22 | 0.50 | 0.82 | 4.10 |
| *GSnet* | 1.22 | 0.63 | 0.74 | 0.94 | 1.11 | 0.11 |
| *aLCnet* | 0.94 | 0.72 | 1.15 | 0.50 | 1.08 | 1.03 |

**Table S6. Computational cost of different methods for predicting global molecular properties via GSnet.** HYDROPRO estimates hydrodynamic properties $R_g$, $R_h$, $D_t$, $D_r$, and $V$. APBS estimates electrostatic solvation free energies $\Delta G_{sol}$. GSnet estimates all properties at once. Structures were obtained from the AlphaFold Protein Structure Database[9], and UNIPROT codes are provided in the table. For GSnet, "Program" represents the total time it takes the program to run, including loading packages, loading models, and converting data, while "Forward" represents the time it takes for a forward pass of the model, which is more relevant for high-throughput predictions. Timings were obtained via the 'time' command in Linux, and via the time module in Python for forward passes of GSnet.

| | **UNIPROT** | **Residues** | **HYDROPRO** | **APBS** | **GSnet[a]** | |
|---|---|---|---|---|---|---|
| | | | | | Program | Forward |
| **Wall Clock Time [s]** | P80967 | 50 | 6.0 | 196.8 | 3.25 | 1.09 |
| | Q16878 | 200 | 25.0 | 469.9 | 3.46 | 1.11 |
| | P80404 | 500 | 244.4 | 2712.8 | 3.92 | 1.17 |
| **User CPU Time [s]** | P80967 | 50 | 69.1 | 188.4 | 19.83 | -- |
| | Q16878 | 200 | 354.4 | 459.9 | 26.17 | -- |
| | P80404 | 500 | 4110 | 2687.3 | 42.72 | -- |
| **Peak memory usage [GB]** | P80967 | 50 | 0.11 | 11.17 | 0.478 | -- |
| | Q16878 | 200 | 0.33 | 12.20 | 0.532 | -- |
| | P80404 | 500 | 1.73 | 30.80 | 0.670 | -- |

[a] Statistics averaged over 10 runs.
